# Disentangling tonic and phasic dopamine in fiber photometry from single recordings to cohorts

**DOI:** 10.64898/2026.11.16.767032

**Authors:** Gabriel Peytral, Zinnera Tariq, Amelie Essmann, Zelai M. Garçon-Poca, David Garcia Ortiz, Caitlin Crook, Jordi Bonaventura, Jordi Abante

## Abstract

Fiber photometry signals combine fast neurotransmitter transients with slower changes in baseline. However, current pipelines separate the two before measuring either, so errors in one carry over into the other. Here we present PhotoSVI, a multi-scale statistical model that splits a recording into a slow tonic signal and discrete release events, estimated together. For each recording, PhotoSVI returns event probabilities over time, the drug effect on event rate with its uncertainty, and the time course of the tonic response, and it pools recordings across a cohort. In simulations matched to experimental recordings, PhotoSVI detected transients and recovered the shape of the tonic response more accurately than existing methods. In mouse dopamine recordings, it revealed an effect that count-based analyses largely missed: a nociceptin receptor agonist enhances the phasic response to morphine while reducing the tonic one, consistent with lower tonic dopamine relieving phasic release from autoreceptor inhibition.

---

Fiber photometry records the bulk activity of a genetically encoded fluorescent sensor [1–4] in a targeted brain region, and has become a standard way to follow neural activity or neurochemical dynamics in behaving animals [5–8]. By enabling the measurement of neural dynamics during behavior, this approach can help link neurochemical changes to the effects of pharmacological interventions. Applied to dopamine, for example, it can address a clinically relevant question: some opioids (such as morphine or fentanyl) act on the same *µ*-opioid receptor yet differ sharply in their behavioral effects and in their overdose risk [9, 10]. If these drugs have distinguishable effects on the dopamine signal, measuring those effects quantitatively would help explain the difference. However, dopamine signaling operates on two time-scales [11, 12]. Fast *phasic* transients, sub-second events tied to discrete occurrences, ride on slow *tonic* shifts in extracellular concentration over minutes. The two are dissociable and support different aspects of behavior [13], and a pharmacological manipulation may act on either or both: opioids reach dopamine neurons by disinhibition [14] and can raise the baseline, change how often transients occur, change how large they are, or any combination. A summary statistic computed over a whole trace collapses these into one number and loses the distinction between them. A practical alternative is to summarize the signal within successive time windows, preserving some temporal structure. We have used this approach previously [10, 15], but it remains sensitive to the choice of window size and alignment and the resulting averages still conflate tonic shifts with changes in phasic activity. Nor do window-based summaries resolve the subjectivity of peak detection [16], which depends on the criteria used to identify discrete transients.

Current tools do not separate tonic from phasic activity. A widely used tool, GuPPy [17], bundles the field’s standard descriptive workflow: motion correction by isosbestic regression, conversion to Δ*F/F* and a z-score, threshold-based transient detection, and behavior-locked peri-event histograms. It is mature and broadly adopted, but every stage rests on point estimates and events are threshold crossings on a signal that still entangles tonic and phasic content. Commercial packages are also in use [18], but their detection rules and defaults are not published, so they cannot be reproduced or benchmarked; we therefore compare against GuPPy, whose procedure is documented. Recent work has added statistical rigor to the downstream question of whether and when a covariate is associated with the signal, modeling trial-level traces as continuous functions of time with functional linear mixed models and reporting joint confidence intervals that respect the trial-within-animal structure [19]. That framework quantifies uncertainty properly, but it remains descriptive: there is no generative account of the signal and no structural separation of baseline from transient. Wallace et al. [16] come closest to the present question, analyzing pharmacologically manipulated dopamine signals and showing that z-scoring against a sliding baseline window can hide the pharmacological effects under study. Their instrument, however, is still a set of deterministic peak detectors returning a count, an amplitude and a width, with no event posterior behind them.

Generative latent-variable modeling has begun to enter the field [20], fitting hidden semi-Markov models and linear dynamical systems to multifiber recordings to recover latent states. Its latent object, however, is a behavioral state rather than the release events themselves, and it offers no tonic–phasic decomposition. The modeling approach we build on comes instead from single-cell two-photon calcium imaging, where spikes are treated as a marked point process and their times and amplitudes recovered by variational Bayesian inference [21]. This is structurally the family used here; it is well established for calcium imaging but has not been applied to bulk photometry.

Here, we present Photometry Stochastic Variational Inference (PhotoSVI), a probabilistic framework based on a core generative model that characterizes dopamine transients and the tonic component together. Because each component maps onto a distinct biological process, the fitted parameters are directly interpretable and comparable across drugs: the tonic curve describes how the drug shifts ambient dopamine, with a rise and a decay set by release and clearance rate constants, while the event intensity and marks are how often transients occur and how large they are. We implement PhotoSVI using probabilistic programming [22] and variational inference [23], which allows the same model to be applied from single recordings to whole studies.

We first introduce a single-trace variant of PhotoSVI and show that it detects phasic events more accurately than the threshold-based detection of standard pipelines [17, 24], while estimating the tonic component from the same fit. We then extend it to a hierarchical model that pools recordings across animals and sessions, follows the structure of the experimental design, in which recordings are nested within animals [25–27], and returns the contrasts of interest with their uncertainty, and we compare it in simulated cohorts with two-stage analyses that test per-recording estimates. Finally, we apply PhotoSVI to a pharmacological question. Morphine increases extracellular dopamine in the nucleus accumbens [28], which contributes to its reinforcing and behavioral effects [10, 29]. Activation of the nociceptin/orphanin FQ peptide (NOP) receptor can modulate these effects and may attenuate morphine-induced dopamine release [30]. Because dopamine signaling contributes to opioid reinforcement, understanding how NOP receptor activation modifies morphine-induced dopamine responses may help clarify its potential in the treatment of opioid use disorder. This interaction also provides a useful test case for a method designed to distinguish changes in tonic dopamine levels from changes in phasic activity. We analyze dopamine recordings from the nucleus accumbens of mice that received morphine after pretreatment with a selective NOP receptor agonist or with vehicle, and ask whether the pretreatment changes the tonic response to morphine, the rate of phasic transients, or both.

## Results

### A generative model for single-trace multi-scale analysis

PhotoSVI is built on a generative model of fiber photometry recordings and of their response to treatment. Although we apply it to dopamine, the model makes no assumption specific to one neurotransmitter, and in principle applies to any genetically encoded sensor whose signal combines a slow baseline with sparse transients, including sensors of other neurotransmitters and neuromodulators and calcium indicators of population activity. Since the tonic and phasic dynamics are entangled in the observed signal, we write both into a single generative model and estimate them together (Fig. 1). A recording is a smooth tonic carrier *x*_*t*_, plus a phasic component *c*_*t*_, plus correlated noise (Methods),

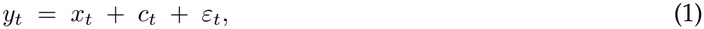

with *ε* autoregressive of order *p*, accounting for the noise structure in the real data (Eq. (8)). Fig. 1a is one 40-minute recording with the fitted carrier in orange: it rises after the injection of an opioid, for example, decays over the remainder, and the transients ride on it throughout. Fig. 1b is the shaded window at the transient scale, where the train of phasic events can be seen. In the proposed model, the phasic component of the signal is a marked point process convolved with a fixed sensor kernel (Fig. 1c). At each time-step a single phasic event *s*_*t*_ can occur, and each event carries an amplitude *a*_*t*_. The product *u*_*t*_ = *s*_*t*_*a*_*t*_ is what the kernel acts on (Methods),

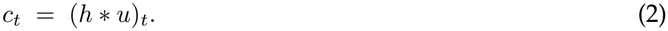

**Figure 1.**
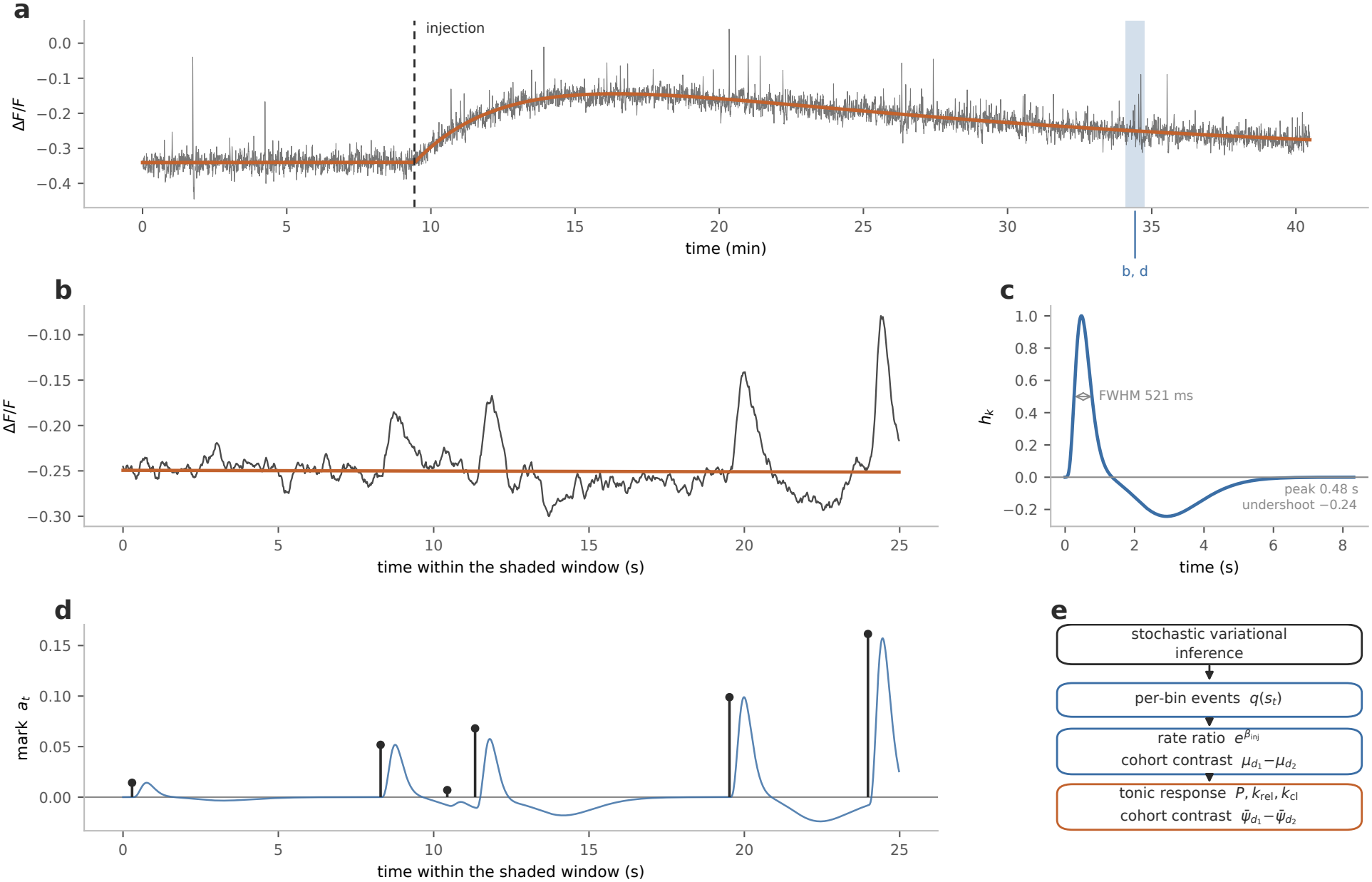
An illustrative example of the model. **a**, A 40-min simulated recording. The tonic carrier (orange) rises after the injection and decays over the rest of the recording, and transients occur throughout. The shaded window is shown in b and d. **b**, The same trace at the time-scale of the transients. **c**, The sensor kernel, with its full width at half maximum, time to peak and undershoot depth. Its taps sum to zero, so the phasic component does not contribute to the signal mean. **d**, The latent process of the model: a sparse train of events with their amplitudes (stems), and the phasic component obtained by convolving it with the kernel (blue). **e**, Event probabilities, the rate ratio and the tonic response, and their contrasts between conditions in a cohort, are read from the same posterior. The recording is simulated at the medium SNR of the benchmark (Methods), and the event train in d is the true one.

Fig. 1d is the latent process the model estimates: a sparse train of events with their amplitudes as stems, and the phasic component they produce once convolved. The rate at which these events occur is determined by the following intensity function:

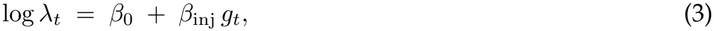

where *g*_*t*_ is 0 before the injection and 1 after it, and we write ***β*** = (*β*_0_, *β*_inj_)^*⊤*^ for the two coefficients. The rate is turned into events through a Bernoulli per bin with 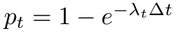. As a result, 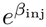 can be read directly as a rate ratio (Eq. (10)).

The tonic carrier has its own parametric form, a Bateman curve [31], given by:

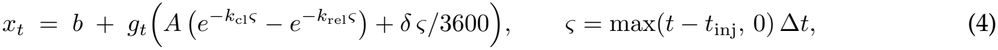

where *b* is the baseline level, *δ* a linear drift per hour, *ς* the time since the injection, *A* the amplitude of the drug response, and *k*_rel_ *> k*_cl_ the release and clearance rate constants. These parameters are estimated jointly with the rest of the model (Eq. (12)). Both quantities are read from a single posterior (Fig. 1e): the per-bin event probabilities *q*(*s*_*t*_), used for detection, and the posterior of *β*_inj_, which gives the rate ratio with its uncertainty.

To estimate the parameters of the model we resort to variational inference, using the following variational approximation (Methods):

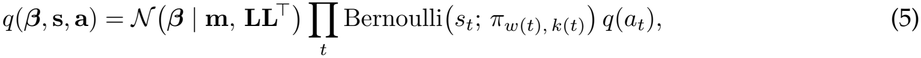

where ***β*** has a Gaussian with mean **m** and free covariance **LL**^*⊤*^, and the events and their marks have one factor per bin. The recording is divided into windows of *W* consecutive bins, bin *t* being position *k*(*t*) of window *w*(*t*), and *π*_*w*(*t*), *k*(*t*)_ is the probability that the onset of that window falls at that position. Each window has a single softmax over *W* + 1 outcomes, no onset or an onset at one of its *W* positions, so the event probabilities of its bins sum to at most one (Eq. (14)). We implemented the model as a module within PhotoSVI in Pyro, a probabilistic programming framework for Python [22].

### Single-trace phasic event detection

One of the main tasks that PhotoSVI can perform is phasic event detection. We compared its detection performance with that of GuPPy, a widely used tool for transient detection in fiber photometry, and of z-peaks, a prominence rule on the z-scored trace used in previous pharmacological studies [10, 15], on simulated recordings of the same length and sampling rate as the experimental ones (Methods). We considered three levels of noise, giving a high, medium and low signal-to-noise ratio (SNR), and matched the power spectrum of the simulated traces to that of the experimental recordings (Supplementary Fig. 1). GuPPy and z-peaks ran at their published default settings and were not tuned on these traces, and every method was evaluated over its full range of thresholds (Methods). PhotoSVI scores events by their posterior probability, so its threshold is a probability, on the same scale for every recording, rather than a level in the units of the signal that depends on its noise.

PhotoSVI detected transients more accurately than both methods at every SNR level (Table 1). Fig. 2 shows a representative window of a recording at medium SNR, with the true event onsets and the events called by each method above the trace. *F*_1_ was 0.783, 0.676 and 0.585 for PhotoSVI at high, medium and low SNR, against 0.669, 0.559 and 0.476 for GuPPy and 0.663, 0.568 and 0.494 for z-peaks. At 90% precision and medium SNR, PhotoSVI recovered 0.508 of the transients, against 0.315 and 0.316. The same fit estimates the tonic response, and PhotoSVI recovered its shape more accurately than the methods in current use (Fig. 2b,c). At medium SNR it overestimated the height of the tonic peak by +1%, against +15% for the low-pass filter of previous studies [10, 15] and +6% for an asymmetric-least-squares baseline, both of which also follow the transients (Methods), and it located the peak more precisely (error +0.3 *±*0.1 against +0.5 *±*1.2 min for the low-pass filter). The area under the tonic response, to which the transients add no net contribution in these simulations, was recovered by all three methods to within a few percent (*−*1.2%, *−*0.1% and *−*2.3%).

**Table 1.** Detection accuracy on simulated recordings. Mean *±* s.d. over 30 simulated recordings per SNR level. SNR is the median event amplitude divided by the standard deviation of the noise (Methods). *F*_1_ is reported at the threshold that maximizes it for each method, and recall at a precision of 90%. PR-AUC, area under the precision–recall curve. Bold, best method for each metric and SNR level.

| SNR | method | *F*<sub>1</sub> | PR-AUC | recall at 90% prec. |
|---|---|---|---|---|
| High (2.9) | GuPPy | 0.669 ± 0.016 | 0.622 ± 0.018 | 0.490 ± 0.033 |
|  | z-peaks | 0.663 ± 0.016 | 0.636 ± 0.023 | 0.484 ± 0.025 |
|  | PhotoSVI | **0.783** ± 0.013 | **0.821** ± 0.017 | **0.685** ± 0.023 |
| Medium (2.0) | GuPPy | 0.559 ± 0.019 | 0.497 ± 0.020 | 0.315 ± 0.030 |
|  | z-peaks | 0.568 ± 0.014 | 0.533 ± 0.019 | 0.316 ± 0.021 |
|  | PhotoSVI | **0.676** ± 0.017 | **0.716** ± 0.016 | **0.508** ± 0.030 |
| Low (1.5) | GuPPy | 0.476 ± 0.020 | 0.394 ± 0.021 | 0.200 ± 0.026 |
|  | z-peaks | 0.494 ± 0.014 | 0.441 ± 0.018 | 0.202 ± 0.031 |
|  | PhotoSVI | **0.585** ± 0.021 | **0.622** ± 0.020 | **0.365** ± 0.033 |

**Figure 2.**
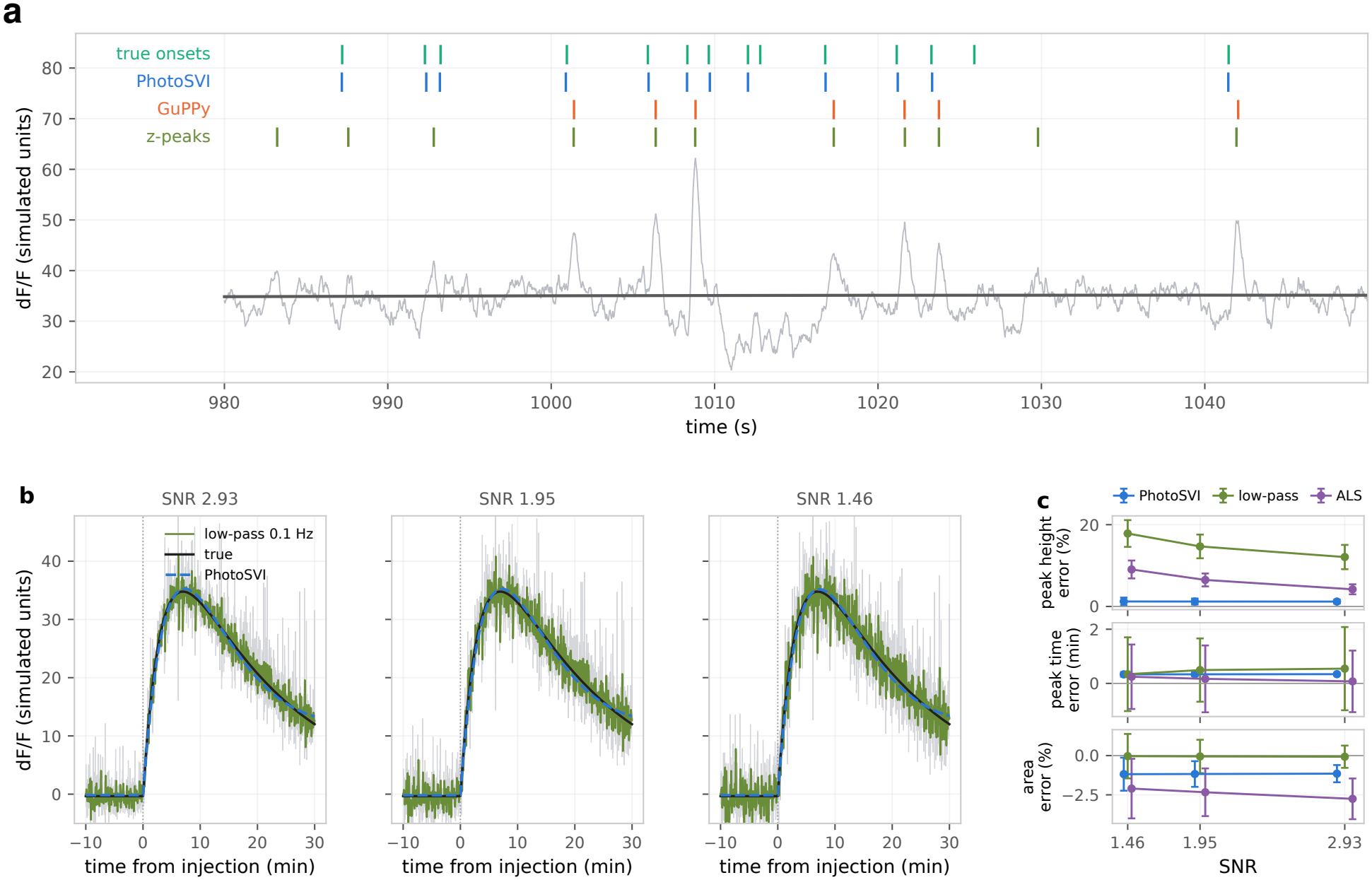
Phasic events and tonic response on simulated recordings. **a**, A window of a recording at medium SNR (2.0; *σ*_*y*_ = 3) with the fitted tonic carrier (dark grey). Above the trace, from top to bottom: true event onsets and the events called by PhotoSVI, GuPPy and z-peaks, each method at the threshold that maximizes its own *F*_1_ on this recording, as in Table 1. **b**, The tonic response of one recording at each SNR level: the simulated truth (black), the tonic carrier fitted by PhotoSVI (blue, dashed) and the 0.1-Hz low-pass filter used in previous studies (green), over 1-s means of the trace (grey). **c**, Relative error in the height of the peak of the tonic response (top), error in its time (middle) and relative error in the area under the tonic response after the injection (bottom) for PhotoSVI, the low-pass filter and an asymmetric-least-squares baseline (ALS), against the SNR; mean *±* s.d. over 30 recordings per level (Methods).

### An extension for cohort-level analysis

A drug effect is a property of a population rather than of a single recording, and averaging per-session estimates discards the between-animal variability that its uncertainty should include. We therefore extended the model to fit all recordings of a study jointly. Each session *j* is a recording of animal *a*(*j*) under condition *d*(*j*), and its two intensity coefficients are written as population terms plus zero-mean random effects (Methods),

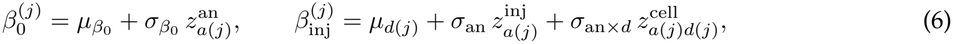

where 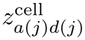 is the effect of the animal-by-condition cell to which the session belongs and every *z* is standard normal (Eqs. (16) and (17)). Each session keeps its own intensity, log 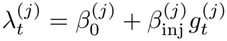, and its own observation model (Eq. (18)), so the single-trace model is the special case of one animal and one session (Supplementary Fig. 2). The effect of condition *d* is the population mean *µ*_*d*_, and the contrast between two conditions, 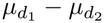, is estimated with the variance components inside its credible interval.

The tonic carrier is pooled in the same way. Its amplitude and rate constants take a mean for each condition and a deviation for each session,

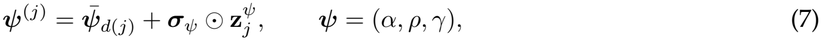

so that the release and clearance rates are estimated for each drug across animals (Eq. (19)), a quantity that a per-session fit does not provide. The baseline level and drift of the carrier remain per session, because they reflect bleaching and fiber placement rather than the drug.

When every animal receives every condition, the animal effect on *β*_inj_ enters both terms of a within-animal contrast and cancels, and the width of the contrast is set by the animal-by-condition interaction *σ*_an*×d*_. When each animal receives a single condition, the contrast compares groups of animals and the between-animal spread *σ*_an_ enters it directly, and PhotoSVI fits only the variance components that the design identifies (Methods). Fitting the hierarchical model also detects the phasic events of every recording, with the event rate of each recording informed by the others through the shared parameters.

### Benchmarking the cohort analysis on simulated cohorts

We compared the proposed hierarchical model with the two-stage analyses that are standard for cohort data, in which each recording is reduced to a single estimate and the estimates are then tested across recordings, unpaired, paired within animal or by a repeated-measures ANOVA. Each two-stage analysis was applied to three kinds of per-session estimate: log rate ratios from the single-trace model, and log rate ratios computed from the transient counts of GuPPy and of z-peaks. Each simulated cohort comprised 5 animals, each recorded once under a control condition and once under treatment, at the medium SNR of the single-trace benchmark (2.0). Treatment increased the log transient rate by Δ relative to control. All analyses were applied to the same simulated cohorts, and the decision threshold of each was calibrated on its own null cohorts (Δ = 0) to a 5% false-positive rate, so that differences in power do not reflect differences in specificity (Methods).

The hierarchical model detected an effect of Δ = 0.50 in 68% of the cohorts, more often than the two-stage analyses of PhotoSVI estimates tested paired within animal (50%), by a repeated-measures ANOVA (50%) or unpaired (54%), whereas the analyses of GuPPy and z-peaks counts detected it in at most 8% and 16% of the cohorts (Fig. 3a). The hierarchical model also estimated the effect with less bias: it recovered 0.76 of the simulated effect at Δ = 0.50, against 0.73 for the paired two-stage analysis, with a smaller root-mean-square error (0.19 against 0.22), and its bias at Δ = 1.00 was *−*0.17 against *−*0.21. The analyses of GuPPy and z-peaks counts recovered 0.05 and 0.05 of the effect (Supplementary Table 1), because their false detections occur at a similar rate before and after the injection and dilute the rate ratio. The 90% intervals of the hierarchical model contained the simulated effect in 80% of the cohorts at Δ = 0.50 and zero in 88% of the null cohorts. Their width matched the spread of the estimate across cohorts (ratio 1.00), so the shortfall reflects the downward bias of the estimate rather than an interval that is too narrow. At low and high SNR (Supplementary Fig. 3 and Supplementary Tables 2 and 3), the hierarchical model and the two-stage analyses of PhotoSVI estimates detected the effect far more often than the analyses of counts, and at low SNR the hierarchical model estimated it with less bias and a smaller error than the two-stage analyses. Across the 250 simulated cohorts, the hierarchical model recovered the tonic response of each condition with median relative errors of 1% in the time and 1% in the height of its peak. For the tonic response, both approaches estimated the contrast between conditions without bias, although the two-stage estimates varied less across cohorts. The low-pass filter and the asymmetric-least-squares baseline also estimated the tonic contrast with little bias, because their overestimate of the tonic peak is common to both conditions, although the bias of the low-pass contrast grew with the effect on the transients (+1% at Δ = 1), and they located the tonic peak about 3 times less precisely than PhotoSVI (Supplementary Fig. 4). The hierarchical model therefore adds power for moderate effects on the transient rate in designs like ours, whereas the two-stage analysis of PhotoSVI estimates remains a sound alternative, in particular at low and high SNR and for the tonic response.

**Figure 3.**
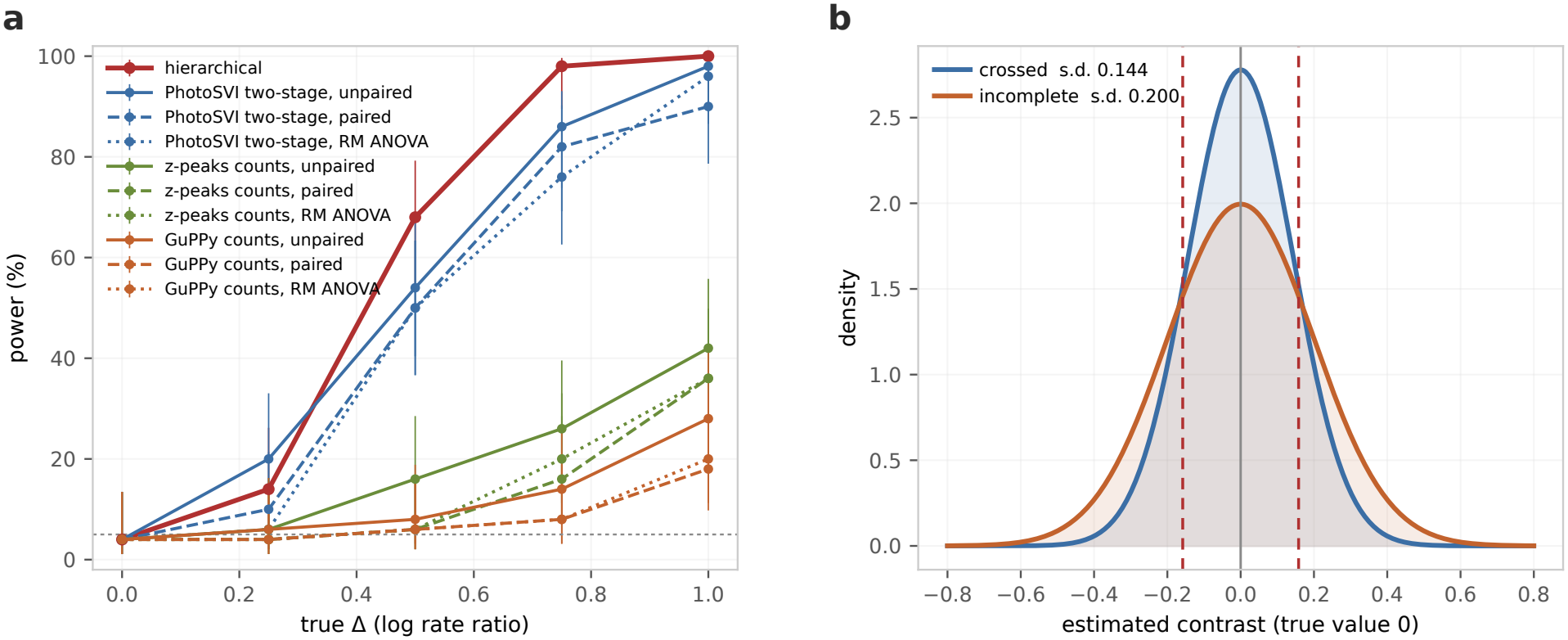
Power to detect a drug effect and the effect of the design. Simulated cohorts at medium SNR (2.0). **a**, Percentage of 50 cohorts per effect size in which each analysis detected the effect, at a false-positive rate calibrated to 5% on its own null cohorts (dotted line); bars, 95% Wilson intervals. Each cohort comprised 5 animals recorded once under each condition; Δ, difference in log event rate between treated and vehicle. Two-stage analyses test per-recording estimates from the single-trace model or from GuPPy or z-peaks event counts, unpaired, paired within animal or by a repeated-measures ANOVA (Methods). **b**, Normal densities with the standard deviation of the estimated contrast across null cohorts (Δ = 0; 50 per design) in the crossed design (0.144; 5 animals recorded under both conditions) and the incomplete design (0.200; 10 animals recorded under one condition each), with the same number of recordings; dashed lines, *±* the standard deviation of an unbiased estimator in the incomplete design (0.158).

We then compared the results obtained by PhotoSVI between a crossed design and an incomplete one in which each animal is recorded under a single condition, with the same number of recordings (10 animals). In both designs the hierarchical model shrank the per-session estimates toward the mean of their condition (Supplementary Fig. 5), as expected from partial pooling: the population distribution of each condition acts as a prior on the effects of its sessions, so that noisy per-session estimates are pulled toward the condition mean. The estimated contrast varied more across null cohorts in the incomplete design (standard deviation 0.200 against 0.144; Fig. 3b), and its posterior was 1.4 times wider. The crossed design therefore estimates the drug effect more precisely, as expected for a design that compares each animal with itself.

### NOP agonism and the dopamine response to morphine

We applied the hierarchical model to a cohort of 7 animals, each recorded in two sessions that ended with the same morphine injection and differed in the pretreatment that preceded it, the nociceptin receptor agonist RO64-6198 or vehicle (Methods). One vehicle session was excluded because of a technical problem, leaving 13 recordings. As a result, the design was crossed for all animals but one, which contributed a single session; the hierarchical model uses the within-animal contrast where it is available. We analyzed the 10 min before the morphine injection and the 30.3 min after it, excluding the handling period of the injection from the likelihood (Fig. 4a). The sensor kernel (Eq. (11)) was the one fitted to these recordings and the noise of each session was estimated from its pre-injection segment (Methods).

**Figure 4.**
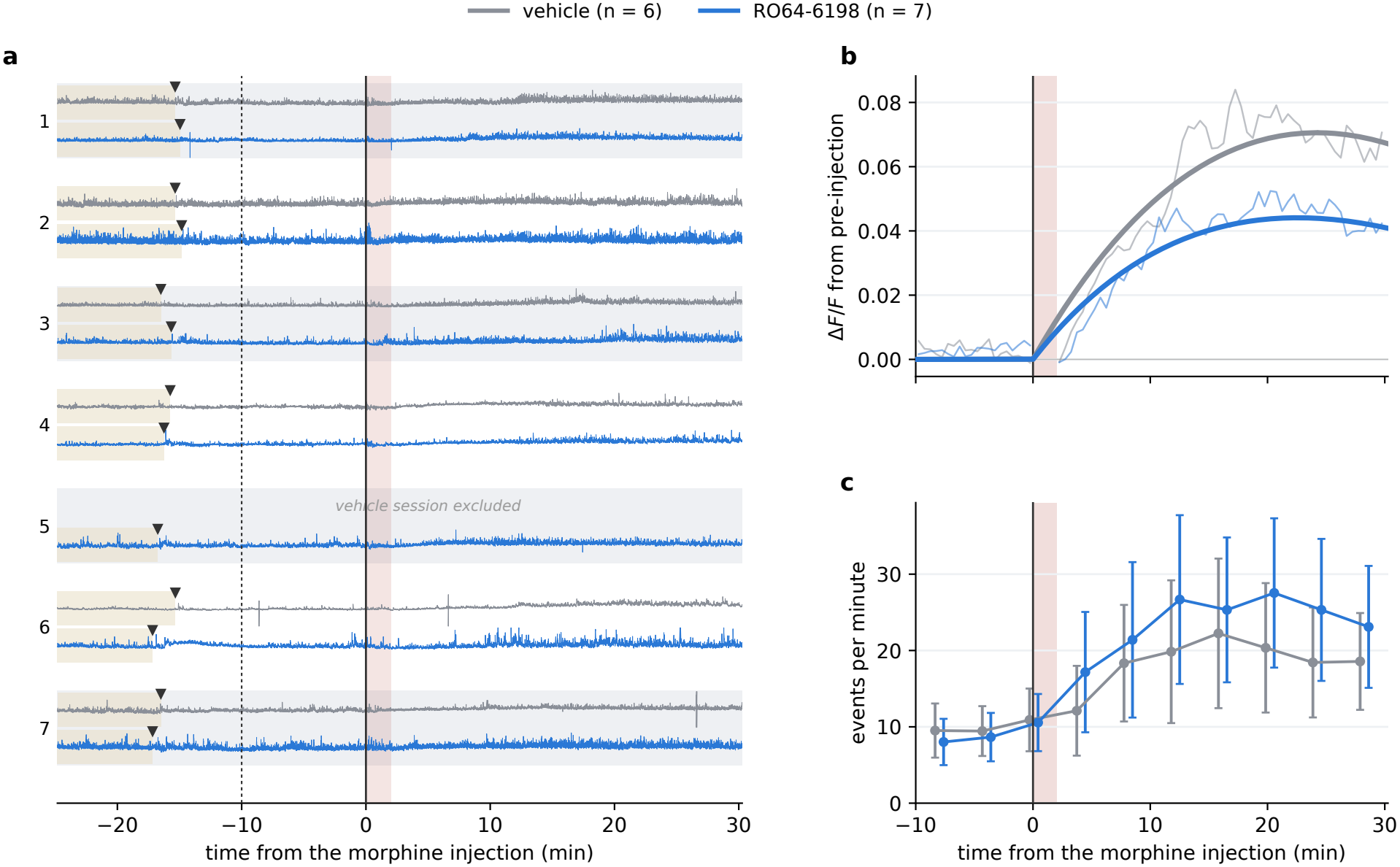
The opioid cohort at both time-scales. Grey, vehicle; blue, RO64-6198; *n*, sessions per condition. **a**, Each complete recording, in minutes from the morphine injection, robustly standardized per session and grouped by animal, with the vehicle-pretreated session above the RO64-6198-pretreated one; the vehicle session of one animal was excluded (Methods). Beige, baseline before the pretreatment injection (triangles); dotted line, start of the analyzed window shown in b and c; shading, window excluded from the likelihood (the 120 s after the morphine injection). **b**, Mean Δ*F/F* of each condition over the analyzed window (thin lines; 30-s bins, each session referenced to its pre-injection level) and the mean tonic component inferred for the same sessions (thick lines). In **b** and **c**, shading marks the same excluded window. **c**, Event rate in 10 time bins, as the posterior expected number of events per minute; mean *±*s.e.m. over the sessions of each condition, with the handling window excluded.

The effect of morphine on transient rate differed between pretreatments: the morphine-induced increase was 1.57 (90% interval [1.13, 2.17]) times larger after RO64-6198 than after vehicle (log-rate contrast: +0.45, Fig. 4c). This difference was detected only within animal: the session means of the two conditions overlap (Fig. 4c), and an unpaired comparison of the per-session estimates did not detect it (*p* = 0.23), whereas a paired two-stage analysis of the same estimates agreed with the hierarchical model (ratio 1.51, 90% interval [1.28, 1.78]). The same paired analysis of GuPPy and z-peaks counts gave smaller ratios, 1.11 (*p* = 0.220) and 1.26 (*p* = 0.055; Supplementary Table 4). The height and the half-height width of the detected transients did not differ between pretreatments (ratios 1.07 and 0.80, 90% intervals [0.85, 1.34] and [0.60, 1.05]; Methods), so the effect of the pretreatment is on the frequency of transients rather than on their shape. The tonic response to morphine, in contrast, was smaller after RO64-6198 (Fig. 4b), with a ratio of fitted peak heights of 0.41 (90% interval [0.40, 0.43]).

## Discussion

Fiber photometry recordings of neurotransmitter dynamics may contain two processes operating on different time-scales: fast transients associated with synchronized discrete release events and slower fluctuations in the signal that may reflect sparse or tonic activity or changes in extracellular neurotransmitter levels. These components can be particularly difficult to disentangle in freely behaving animals, where transients are not necessarily time-locked to identifiable behavioral events, or in pharmacological experiments, where drug administration can introduce sustained changes in the signal over the course of a recording. Current analysis pipelines separate the two before measuring either, first estimating a baseline and then counting the transients that exceed a threshold above it. PhotoSVI estimates both jointly with a generative model in which the transients are a marked point process whose rate can vary with the experimental condition, slower changes in the signal are captured by a time-varying baseline, and residual fluctuations are modeled as an autoregressive process. Fitted by variational inference, the model returns the probability of an event in every bin, the change in event rate caused by the drug with its uncertainty, and the parameters of the tonic response, all from a single posterior. Its hierarchical extension pools these quantities across the animals and conditions of a cohort. Because a single model is fitted, the phasic and tonic estimates are consistent with each other: transients are detected against the same baseline that defines the tonic response, rather than against a baseline fixed by an earlier step, as in calcium imaging, where baseline and activity are estimated jointly [32].

On simulated recordings matched to experimental data in their kernel, noise spectrum and signal-to-noise ratio, PhotoSVI detected transients more accurately than GuPPy and than the prominence rule used in previous pharmacological studies at every noise level, with an *F*_1_ score 17% to 19% higher than the better of the two. On the same recordings it recovered the shape of the tonic response more accurately than a low-pass filter or an asymmetric-least-squares baseline, which overestimate its peak because they also follow the transients. Detection is, however, an intermediate step: most pharmacological photometry experiments ask by how much a drug changes the frequency of transients, and PhotoSVI estimates this rate directly as a model parameter, with an interval. On simulated cohorts at the SNR of the experimental recordings, the hierarchical model detected a drug effect more often than the best two-stage analysis of PhotoSVI estimates and estimated it with less bias, particularly for large effects, whereas traditional analyses of event counts recovered only a small fraction of the effect. In addition, the advantage of this model grows as the per-session estimates become noisier. The cohort model also recovered the tonic response of each condition, which a count-based analysis does not estimate, with median errors of 1% in the time and 1% in the height of its peak.

PhotoSVI is also a simulator with known ground truth at every level, which could be used to compare other detection tools on equal terms and along axes that experimental data cannot vary on demand, such as noise level, effect size and cohort design. Run with the variance components estimated from a pilot cohort, the simulator gives the expected power of a planned study for a given number of animals and design, which helps to avoid the low statistical power behind many irreproducible findings in neuroscience [33, 34]. A two-stage analysis cannot provide this, because it does not estimate those components. In our simulations, for example, a crossed design with 5 animals estimated the drug effect more precisely than an incomplete design with 10 animals and the same number of recordings.

In the opioid cohort, consisting of 13 recordings from 7 animals, the hierarchical model estimated that the increase in transient rate caused by morphine was 1.57 times larger after pretreatment with a nociceptin receptor agonist than after vehicle, whereas the tonic response to morphine was smaller. The analyses of transient counts used in previous studies would have underestimated this effect: paired within animal, GuPPy and z-peaks counts gave ratios of 1.11 and 1.26, consistent with the attenuation of the effect by count-based analyses in the simulated cohorts.

Microdialysis, which samples extracellular dopamine over minutes and cannot resolve individual transients, has shown that nociceptin reduces the morphine-induced increase in dopamine, particularly in the shell of the nucleus accumbens (NAc), without altering basal dopamine [35, 36], consistent with the smaller tonic response observed here. An opposite effect on the rate of transients is less expected and, to our knowledge, has not been described; a measure of mean concentration could not have revealed it. Two mechanisms could contribute to it. First, in the tonic–phasic framework proposed by Grace [11], tonic extracellular dopamine sets the background stimulation of dopamine receptors, including autoreceptors, so a lower tonic level could leave transients less constrained. Second, nociceptin hyperpolarizes both dopamine and non-dopamine (putative GABAergic) neurons in the ventral tegmental area (VTA) but also reduces spontaneous and evoked GABAergic inhibitory postsynaptic potentials in dopamine neurons [37], a partial disinhibition that would oppose its direct inhibitory action on these cells. Morphine itself acts by disinhibition, increasing dopamine neuron activity through *µ*-opioid (MOP) receptor-mediated inhibition of VTA GABAergic neurons [14]. NOP activation also inhibits NAc-projecting VTA neurons while having little effect on evoked dopamine release at NAc terminals ex vivo [38], consistent with compartment- and projection-dependent actions. The opposing changes in transient frequency and tonic signaling could therefore reflect distinct NOP-sensitive mechanisms within mesolimbic circuitry, whose relative contributions remain to be established.

This dissociation may also bear on the effects of NOP activation on opioid reward. Nociceptin prevents morphine-induced conditioned place preference without affecting the development of locomotor sensitization [39], and RO64-6198 blocks the acquisition and reinstatement, but not the expression or extinction, of morphine conditioned place preference [40]; NOP activation therefore interferes with some aspects of opioid reward rather than suppressing opioid-induced behavior uniformly. A smaller tonic dopamine response could contribute to the attenuation of the sustained motivational effects of morphine, while the increased frequency of transients indicates that dopamine signaling is not suppressed at every time-scale. NOP receptor activation can also contribute to analgesia while counteracting some adverse effects of MOP receptor activation [41], which raises the possibility that differential modulation of dopamine signaling contributes to separating opioid analgesia from reward. Both possibilities remain speculative: neither reinforcement nor analgesia was assessed here, and the pharmacology of dual MOP/NOP agonists cannot be extrapolated directly to the separate administration of RO64-6198 and morphine.

The formulation can be extended in several directions. The log intensity is a linear predictor, so several interventions per recording can be modeled with a coefficient and an injection gate for each, and time-varying covariates such as locomotion or reward delivery can enter as additional terms, which would allow the drug effect to be estimated while controlling for behavior. Discrete behavioral states obtained by unsupervised segmentation of video could enter in the same way, relating the rate of transients to the behavior during which they occur, as has been done for striatal dopamine and behavioral syllables [8]. The drug is currently assumed to change the rate of transients but not their size. However, letting its location change at the injection would estimate an amplitude ratio alongside the rate ratio, separating the frequency of release events from the amount released or cleared per event. Multichannel recordings could share a latent event train observed through channel-specific kernels and noise, turning the isosbestic channel from a preprocessing step into a modeled quantity, and recordings from two regions could be modeled with coupled intensities. Finally, the resolution of closely spaced events is bounded by the averaged transient, whose half-width of about half a second reflects both the kinetics of the sensor and the dissociation and clearance of dopamine; because PhotoSVI calibrates its kernel to the sensor used, faster sensors would translate directly into finer resolution.

In summary, PhotoSVI replaces the sequence of baseline estimation, thresholding and counting with a single probabilistic model of the recording. It detects transients more accurately, estimates the drug effect on their rate with its uncertainty, recovers the tonic response and extends to cohorts. Applied to a cohort of opioid recordings, it shows that pretreatment with a nociceptin receptor agonist enhances the phasic dopamine response to morphine while reducing its tonic response, a dissociation between time-scales that a measure of mean dopamine concentration would not reveal.

## Methods

### Generative model

We model a fiber-photometry trace as the sum of a sparse, fast *phasic* train of dopamine transients and a slow *tonic* carrier under a single probabilistic model. For a recording of *T* bins at spacing Δ*t* with the injection at bin *t*_inj_,

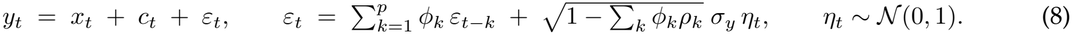

The residual is autoregressive of order *p* [42], with the noise *η*_*t*_ scaled so that *σ*_*y*_ is the marginal standard deviation; *ρ*_*k*_ are the autocorrelations, and *p* = 0 gives white noise. Unlike white noise, we found that second-order autoregressive noise reproduces the spectrum of the experimental recordings (Supplementary Fig. 1), and it improves on first-order noise in the Akaike information criterion in all 10 recordings (by 1,781 to 18,740), whereas higher orders improve it by smaller amounts. The tonic and phasic components are identified by their time-scales, minutes and seconds, respectively.

To account for phasic events of different intensities, the transients form a marked point process [43]. Thus, each event carries an amplitude, which separates the effect of a drug on how often transients occur from its effect on how large they are. First, the conditional intensity of the point process is given by

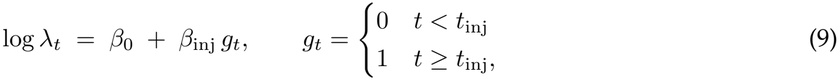

and events are drawn one per bin through the complementary log-log link [44], so that *p*_*t*_ is the probability that a Poisson process of rate *λ*_*t*_ has at least one event in the bin,

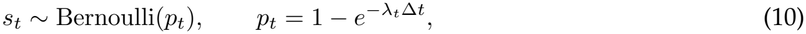

so that *β* is a log rate and 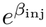 is a rate ratio. Second, each event carries a mark *a*_*t*_ LogNormal(*µ*_*a*_, *σ*^2^), and the marked train *u*_*t*_ = *s*_*t*_*a*_*t*_ is convolved with a fixed causal kernel *h*, which can be adjusted to the specific sensor used (e.g., dLight1.3b [1]), given by

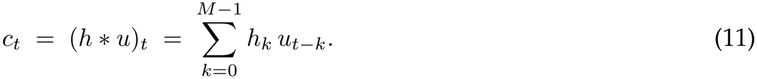

The injection enters through *β*_inj_ alone, while *µ*_*a*_ and *σ*_*a*_ are shared before and after it; the current version of the model therefore represents a drug effect on the frequency of transients but not on their size. Nevertheless, differences in size can also be studied via the posterior distribution of *{a*_*t*_*}*_*t*_.

We assume the sensor kernel has zero area, i.e., ∑_*k*_ *h*_*k*_ = 0, resulting in a fast positive lobe minus a slower lobe scaled so that their areas cancel, so that the phasic component does not contribute to the mean of the signal, carried by the tonic component alone. The kernel families implemented in PhotoSVI are listed in Supplementary Table 5. Here we use a difference of two gamma lobes fitted to the peak-triggered average of isolated transients in the experimental recordings, which describes the kinetics of the dLight1.3b sensor used to acquire them. Because the kernel depends on the sensor, and through release and clearance on the region recorded, PhotoSVI includes a calibration function that extracts isolated transients from a user’s recordings, fits each family to their average and suggests the one with the lowest error on held-out transients, so that recordings made with sensors of other kinetics [2, 3, 45] are modeled with their own kernel.

The tonic carrier is a Bateman curve [31], switched on at the injection, plus a level and a linear drift,

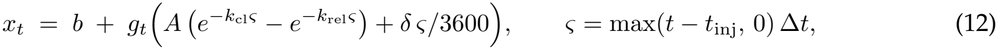

with release and clearance rate constants *k*_rel_ *> k*_cl_ *>* 0. For fitting, the amplitude *A* is replaced by the height *P* = *e*^*α*^ of the response, in the units of the trace, and the rate constants are written as *k*_cl_ = *e*^*ρ*^ and *k*_rel_ = *e*^*ρ*^ + *e*^*γ*^, which enforces their order. The drift *δ* is expressed per hour so that its gradient has a similar scale to those of the other parameters. The five tonic parameters are initialized by least squares and refined jointly with the rest of the model as point estimates.

Writing ***θ*** for the tonic parameters, the joint density of the observations and the latent variables is (Supplementary Fig. 2)

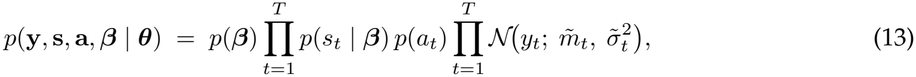

where *m*_*t*_ = *x*_*t*_ +*c*_*t*_, *r*_*t*_ = *y*_*t*_ *−m*_*t*_, and for *t > p* the conditional mean and variance are 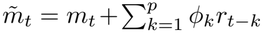 and 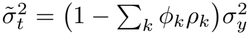.

### Single-trace model training

The kernel and the autoregressive coefficients are fixed, and *σ*_*y*_ is estimated from the successive differences of the pre-injection segment of each recording, corrected for the autocorrelation of the residual, prior to fitting the probabilistic model. Then, PhotoSVI uses the following variational family when analyzing a single trace:

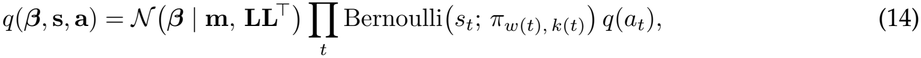

where ***β*** = (*β*_0_, *β*_inj_)^*⊤*^ and **L** is a 2 *×*2 lower-triangular matrix. *β*_0_ and *β*_inj_ have wide Normal priors, centered on a matched-filter estimate of the baseline rate and on zero, respectively (Supplementary Table 6). A single transient produces a correlation lobe several bins wide, and an independent factor per bin would spread fractional event mass across the lobe, inflating the estimated rate. As a result, PhotoSVI divides the recording into windows of *W* = 10 consecutive bins (208 ms); bin *t* is position *k*(*t*) of window *w*(*t*), and each window has ***π***_*w*_ = softmax(**z**_*w*_) over *W* + 1 outcomes, i.e., no onset or an onset at one of its *W* positions. The mark posterior is shared within each window, *q*(*a*_*t*_) = LogNormal 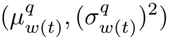. The amplitudes are fixed in units of the noise, with the same distribution for every recording. The tonic parameters are point estimates, and the other fixed settings are listed in Supplementary Table 6.

The evidence lower bound (ELBO), given in this case by

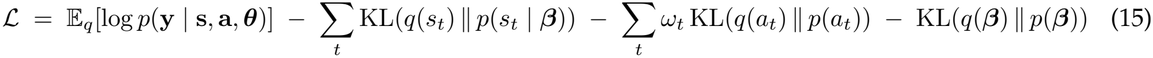

where *ω*_*t*_ = max *π*_*w*(*t*), *k*(*t*)_, *f* weights the divergence of each mark by the event probability of its bin, with a floor *f* = 0.1, is maximized over the variational parameters and parameters ***θ***. The weight follows from the fact that a mark exists only where there is an event, so its divergence counts in proportion to the probability of an event in that bin. To pass gradients through the discrete events, the reconstruction term replaces each event by a binary Concrete variable, a continuous relaxation of the Bernoulli distribution that approaches it as the temperature *τ* goes to zero [46, 47]; we use *τ* = 0.2. Due to this relaxation, bins with a negligible event probability still contribute a little to the reconstruction and can make training unstable; the floor *f* in *ω*_*t*_ prevents this.

Starting values for the tonic parameters, the coefficients and the event probabilities come from a least-squares pre-fit of the tonic carrier and a matched filter (Supplementary Note 1). The model is fitted by stochastic variational inference using Pyro [22]. The bound is maximized with Adam (learning rate 0.01), and the fit stops when the posterior mean of *β*_inj_ has stabilized, by the criterion given in Supplementary Table 6.

### Hierarchical cohort model

The cohort model estimates the drug effect as a population parameter with between-animal variability, rather than averaging per-recording estimates, whose uncertainty does not include the between-animal variability. Sessions are indexed by *j* = 1, …, *S*, each a recording of animal *a*(*j*) under condition *d*(*j*). Thus, an animal may be recorded under one condition or several. The coefficients of session *j* are a population term plus zero-mean random effects,

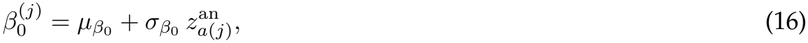

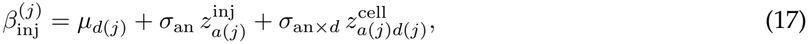

with every *z*_*•*_ *∼ N* (0, 1) and each *σ*_*•*_ log-normal. The intensity of session *j* is

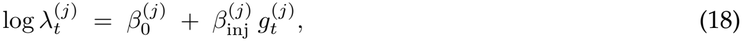

with a per-session gate 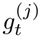 because the injection time differs between recordings. The animal-by-condition interaction requires animals recorded under more than one condition: it is included in the crossed design and omitted in the incomplete one, where it cannot be distinguished from the animal term. Further levels, and the rule by which the levels are chosen from the design, are described in Supplementary Note 2.

We assume that *µ*_*d*_ *∼ N* (0, 1), which gives rate ratios between 0.14 and 7.1 a prior probability of 95%. For the baseline rate we use an empirical Bayes prior, whose center is estimated from the data rather than fixed in advance [48, 49]. This suits parameters whose scale changes between datasets, such as the baseline rate, which depends on the sensor, the region and the recording. Because the data are then used twice, the posterior can be slightly too narrow, but the effect is small when the prior is wide relative to the information in the data. Thus, we assume 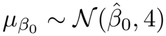, where 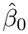 is the mean over sessions of a matched-filter estimate of the baseline rate (Supplementary Note 1). It is wider than the prior on *µ*_*d*_ because the 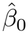 estimate is quite noisy (Supplementary Note 3).

The tonic response is pooled by condition. Its amplitude and rate constants (*α, ρ, γ*), the unconstrained parameters of *P, k*_cl_ and *k*_rel_, have a per-condition mean and a per-session deviation,

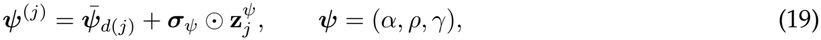

We also use an empirical Bayes prior for the tonic response, 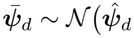,diag(1, 0.25, 0.25), where 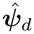 is the mean least-squares pre-fit of the sessions of condition *d*. Because this center differs between conditions, it could favor a tonic contrast. However, we found that in simulated cohorts with the same tonic response in both conditions, the estimated tonic contrast was unbiased (Supplementary Fig. 4). The level *b* and the drift *δ*, which reflect bleaching and fiber placement rather than the drug, are estimated per session without pooling. The per-session deviation is included only when a condition has more than one session.

### Fitting the hierarchical model

Inference follows the single-trace model, with the same relaxation, event divergence, windowed event factor and per-window marks. Each session has its own noise scale 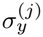, estimated from its pre-injection segment as in the single-trace model. Writing Θ for the population terms 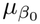, the *µ*_*d*_ and the logarithm of each variance component), **z** for the non-centered effects of Eqs. (16) and (17), and 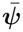 and **z**^*ψ*^ for the per-condition tonic means and per-session tonic deviations of Eq. (19), the variational family is

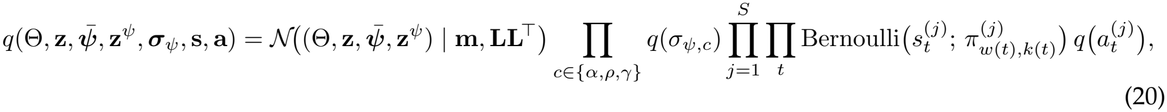

with **L** lower triangular. The first factor is a Gaussian with full covariance over the population terms, the non-centered effects and the tonic means and deviations; the scales of the tonic hierarchy have independent log-normal factors, and the events and marks of each session have the windowed factors of the single-trace model (Eq. (14)). The per-session level and drift are point estimates. We report central 90% posterior intervals throughout, the level at which their coverage was evaluated in the simulated cohorts. An interval that excludes zero corresponds to a one-sided test at the 5% level. Finally, as in the single-trace case, the hierarchical model is trained using Pyro, and the bound is maximized with Adam (learning rate 0.01), with all recordings of a cohort fitted as one batch on a GPU, and the fit stops when the posterior means of the *µ*_*d*_ have stabilized, by the criterion given in Supplementary Table 6.

### Single-trace detection benchmark

Benchmark recordings were drawn from Eqs. (8) to (12) with the length, sampling rate and injection time of the experimental recordings (40 min at 48 Hz, injection at 10 min) and with parameters calibrated on them (Supplementary Note 3), resulting in a power spectrum that closely matches that of the experimental recordings (Supplementary Fig. 1). To consider different noise conditions, we simulated recordings under three different signal-to-noise ratio (SNR) conditions. More precisely, we defined the SNR as the median mark over the noise standard deviation, 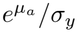 and considered three noise levels, *σ*_*y*_ = 2, 3 and 4, resulting in a high, medium and low SNR of 2.9, 2.0 and 1.5, and we simulated 30 recordings per level. The model fixes the amplitude distribution in units of the noise, which matches the simulated amplitudes at medium SNR. At high and low SNR the simulated amplitudes are larger or smaller than the model assumes, so these two levels also test the model when its amplitude distribution is wrong.

We compared the detection performance of PhotoSVI, GuPPy [17] and the detection rule of our previous pharmacological studies [10, 15, 24]. PhotoSVI was fitted to each recording with the settings of the singletrace model. It scores each bin by its posterior event probability *q*(*s*_*t*_=1) and detects an event wherever this probability peaks above a threshold. GuPPy [17] was reimplemented from its published transient rule and run at its default settings, without tuning. Within each non-overlapping 15-s segment, it computes the median and the median absolute deviation (MAD), excludes samples above the median plus 2 MAD, recomputes both on the remaining samples, and calls every interior local maximum above the median plus 3 MAD a transient, scored by its height above that median. We also evaluated the detection rule used in previous pharmacological photometry studies from one of our laboratories [10, 15, 24], which we refer to as z-peaks: the trace is smoothed with a Savitzky–Golay filter over 0.1 s, z-scored against its pre-injection segment, and every local maximum with a prominence above 1.6 standard deviations is called a transient, scored by its prominence. GuPPy and z-peaks report the peak of a transient and PhotoSVI its onset, so each method’s median offset from the true onsets was subtracted from its detection times before matching (Supplementary Note 4).

We considered a detection to be a true positive if it falls within *±*354 ms of an onset not already matched to a higher-scoring detection. Precision is TP*/*(TP + FP), recall is TP*/*(TP + FN) and *F*_1_ = 2*PR/*(*P* + *R*), where TP, FP and FN are the numbers of true positives, false positives and undetected onsets. Here, we report *F*_1_ at the threshold that maximizes it, the area under the precision–recall curve (PR-AUC), computed as the average precision, and the recall at 90% precision, so that no method is evaluated at its nominal threshold.

### Tonic response estimation

The tonic response estimated by PhotoSVI on the benchmark recordings was compared with the slow component extracted by two methods that do not model the transients. The first is the low-pass filter used in previous pharmacological photometry studies [10, 15]: a fourth-order Butterworth filter at 0.1 Hz, applied causally. The second is an asymmetric-least-squares baseline [50], which penalizes the curvature of the baseline and weights the samples above it less than those below (smoothness 10^6^*× f*_*s*_, asymmetry 0.01, 10 iterations). Both were applied to the trace in its own units; the z-scoring of the original pipeline rescales the curve and the truth alike and leaves relative errors unchanged. Each curve, and the simulated tonic carrier, was summarized by the height of its maximum after the injection above its mean from 5 min to the injection, by the time of that maximum, and by its area above that level from the injection to the end of the recording. Then, we compared the errors against the simulated values over the 30 recordings of each SNR level. In the cohort benchmark, the same two methods were applied to every recording, and the log ratio of peak heights and the difference of peak times between the treated and vehicle sessions of each animal were averaged over animals, as for the two-stage analysis of PhotoSVI’s single-trace estimates.

### Cohort benchmark

The cohorts of Fig. 3 and Supplementary Table 1 were drawn from the hierarchical model with a crossed design: 5 animals, each recorded once under vehicle and once under treatment, with the kernel and noise measured on the recordings and amplitudes simulated as in the single-trace benchmark, at its three SNR levels. The treatment raises the log event rate after the injection by Δ*∈* {0.00, 0.25, 0.50, 0.75, 1.00} relative to vehicle, and the baseline rate and the response vary between animals, with an animal-by-condition interaction and per-session variability in the baseline rate. We simulated 50 cohorts per Δ at medium SNR, reported in the main text, and fewer at low and high SNR for Δ *∈* {0, 0.5, 1} (Supplementary Fig. 3 and Supplementary Tables 2 and 3). All analyses were applied to the same recordings.

The two-stage analyses first estimate a log rate ratio 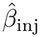 per recording, either as the posterior mean of the single-trace model or from the events called by GuPPy or z-peaks, as log (*n*_post_*/T*_post_)*/*(*n*_pre_*/T*_pre_) with *n* the number of events and *T* the duration before and after the injection. The unpaired test is Welch’s *t*-test of the treated against the vehicle estimates; the paired test is a one-sample *t*-test of the within-animal differences; the repeated-measures ANOVA tests the condition-by-period interaction on the event rates before and after the injection (for the single-trace model, 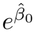 and 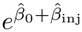, computed as the square of a paired *t* on the per-animal difference of differences, which gives a test but no estimate of Δ. The hierarchical analysis uses |*z*|, the posterior mean of *µ*_treated_ *−µ*_vehicle_ divided by its posterior standard deviation. The single-trace and hierarchical fits use the same settings (Supplementary Table 6).

Power was compared at the same false-positive rate for all analyses. For each analysis, the threshold was the value exceeded by 5% of its statistics on the null cohorts (Δ = 0), and the power at each Δ is the fraction of cohorts above that threshold. The bias and root-mean-square error in Supplementary Table 1 are those of each analysis’s estimate of Δ: the difference between the mean treated and vehicle estimates (unpaired), the mean within-animal difference (paired) or the posterior mean of the contrast (hierarchical).

The design comparison of Fig. 3 uses null cohorts (Δ = 0) at medium SNR with the same number of recordings in two designs: crossed, with 5 animals recorded under both conditions, and incomplete, with 10 animals recorded under one condition each.

### Experimental recordings and preprocessing

Male (n = 2) and female (n = 5) adult wild-type C57BL/6 mice (Charles River Laboratories; 20–30 g, 8– 16 weeks old) were housed under controlled temperature and humidity on a 12-h light–dark cycle with food and water ad libitum. The cohort was too small to compare sexes, and sex was not included as a covariate. Experiments were performed during the light phase. All experiments were approved by the Comitè d’Experimentació Animal of the Generalitat de Catalunya (CEEA40/22 and CEEA167/22) and conducted in accordance with the Guide for the Care and Use of Laboratory Animals (National Institutes of Health, 1996) and European Communities Council Directive 2010/63/EU.

For expression of the dLight1.3b sensor and fiber implantation, mice were anesthetized with isoflurane (4.5% induction, 1.5% maintenance) and placed in a stereotaxic apparatus (RWD Life Sciences). Mice were unilaterally injected with 500 nL of an adeno-associated viral vector encoding dLight1.3b (ssAAV-1/2-hSyn1-chI-dLight1.3b-WPRE-bGHp(A), 7.6 *×*10^12^ vg mL^*−*1^; ETH Viral Vector Facility, University of Zurich) into the nucleus accumbens shell (AP +1.1, ML 0.7 from bregma, DV *−*4.2 from the dura, according to the Paxinos atlas) with a 2-*µ*L microsyringe (Hamilton Neuros) and a syringe pump (Stoelting) at 50 nL min^*−*1^. The syringe was left in place for 5 min. An optic fiber (6 mm, 400-*µ*m core, 0.5 NA; RWD Life Sciences) was then lowered to the nucleus accumbens shell and secured with dental cement (Dentalon, Kulzer).

Recordings were made 3–4 weeks after surgery, to allow recovery and sensor expression, while mice explored a 20*×* 15 cm Plexiglas cage. Each session comprised a 10-min baseline, an intraperitoneal injection of vehicle (saline with 5% DMSO) or of the nociceptin receptor agonist RO64-6198 (1 mg kg^*−*1^ in vehicle), a 15-min recording, a subcutaneous injection of morphine hydrochloride (20 mg kg^*−*1^ in saline) and a 30-min recording. Fluorescence was excited at 470 nm (dopamine-dependent) and 405 nm (isosbestic control for bleaching and movement artifacts) and collected at 500–540 nm with a Fiber Photometry Console and Neuroscience Studio v6 (Doric Lenses). The light sources were controlled with T-Cube LED drivers (Thorlabs), and the LEDs, dichroic mirrors and photomultipliers were integrated in a Doric Minicube. Both channels were acquired simultaneously on the same detector by lock-in amplification at 12,048 Hz, low-pass filtered at 12 Hz and downsampled 251-fold to 48 Hz. Injection times were taken from the events recorded by the acquisition software or from the experimental log; for two vehicle sessions in which neither was recorded, the times of the protocol were used.

The cohort comprised 7 animals, each recorded in two sessions that ended with the same morphine injection and differed in the pretreatment, RO64-6198 in one and vehicle in the other. One vehicle session was excluded because of a technical problem, leaving 13 sessions. The isosbestic channel was smoothed with a 0.5-s moving average, and 1 s was trimmed from each end of both channels. The baseline of the sensor channel was estimated by asymmetric least squares (smoothness 10^6^*× f*_*s*_, asymmetry 0.01, 10 iterations), and the isosbestic channel was mapped onto this baseline by linear regression over the first 10 min of the recording, or up to the morphine injection if earlier. Δ*F/F* was computed as (*F− F*_0_)*/F*_0_, with *F*_0_ the mapped isosbestic channel. The sensor channel was not smoothed, because the transients occupy the band a smoothing filter would remove. The analysis window runs from 10 min before the morphine injection to the end of the recording and was not detrended, because a linear trend fitted across the injection distorts the trace around it; slow trends are modeled by the drift term. The 120 s after the start of the morphine injection, or the recorded handling period if longer (9 to 80 s; 5.0% of the analyzed samples), were excluded from the likelihood, together with the following *p* bins of the autoregressive emission: handling produces transient-shaped artifacts at the time the rate is expected to change, and both channels decrease for 1–2 min after the injection.

Because the model fixes the amplitude distribution and the kernel, the height and width of the transients were compared between pretreatments outside the model. Transients were taken as the local maxima of the posterior event probability above 0.5 with no other such maximum within 1.5 s. Their height was the maximum of the trace minus the fitted tonic component over the following second, relative to the median of the preceding 0.5 s, and their width was the time between the two half-height crossings. For each session, the median after the injection (from 2 min) was divided by the median before it, and the log ratio between pretreatments was averaged over the animals recorded under both, with a *t* interval.

For comparison with the hierarchical model, the effect of the pretreatment on the morphine response was also estimated by two-stage analyses. For each session, the rate of transients after the morphine injection (excluding the masked window) was divided by the rate before it, with rates taken from the posterior expected number of events of the cohort fit or from the calls of GuPPy and z-peaks at their default rules. The log ratios were compared between pretreatments with a paired *t*-test over the animals recorded under both and with an unpaired *t*-test over all sessions (Supplementary Table 4).

## Data availability

The fiber photometry recordings analyzed in this study are available from the corresponding authors upon reasonable request. The simulated recordings can be regenerated with the code below.

## Code availability

PhotoSVI, including the scripts that reproduce the analyses and figures of this study, is available at https://github.com/AbanteLab/photosvi.

## Use of large language models

Large language models were used to assist in writing the text and as support in writing the analysis and figure code. All outputs were reviewed and verified by the authors, who take full responsibility for the content.

## Acknowledgements

This work was supported by grants RYC2023-043080-I (J.A.), PID2025-174934OA-I00 (J.A.), PID2023-147013OB-I00 (J.B.), and CNS2024-154735 (J.B.), all funded by MICIU/AEI/10.13039/501100011033. It was further supported by a Synapse Grant awarded by the Institut de Neurociències, Universitat de Barcelona, funded in part through the María de Maeztu award CEX2021-001159-M, also funded by MI-CIU/AEI/10.13039/501100011033. The funders had no role in study design, data collection and analysis, decision to publish, or preparation of the manuscript.

## Author contributions

Z.T. developed an initial approach based on deep Markov models, including a first model of the tonic response, as part of a master’s thesis supervised by J.A. G.P. implemented the first version of the latent marked point process model under the supervision of J.A. D.G.O. contributed to the processing of the experimental data, under the supervision of J.A. and J.B. A.E., Z.G. and C.C. performed the surgeries and pre-processed and curated the photometry data. J.B. supervised the experimental work and the pre-processing of the experimental data. J.A. developed the final methodology and software, performed the analyses and wrote the manuscript with J.B. J.B. and J.A. designed the study. All authors reviewed and approved the final manuscript.

## Supplementary Notes

### Supplementary Note 1: Initialization of the single-trace model

We initialize the model from a matched filter. First, we fit Eq. (12) by least squares to the trace median-filtered over 10 s, without and then with the drift. If the fit with drift returns a height below 10^*−*5^, we keep the fit without drift. We then cross-correlate the residual of this pre-fit, *r*_*t*_, with the kernel, 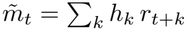, and standardize the result over the pre-injection segment, *m*_*t*_ = (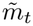 median 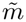)*/* sd 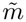 (over the whole trace if that segment is shorter than 250 bins). The Gaussian factor over ***β*** starts at mean 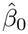 with Cholesky factor **L** = 0.1 **I**, where 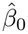 = log max(*n*, 1)*/*(*t*_inj_Δ*t*) and *n* is the number of local maxima of *m*_*t*_ above 2.5 before the injection, after non-maximum suppression over 15 bins. The window scores start at *z*_*w,k*_ = 2 (*m*_*t*_*−* 2.5), and the score of the no-onset outcome is fixed at zero, so that the initial guide behaves like a matched-filter detector at that threshold; a final window shorter than *W* is padded with scores of -30. The mark posterior starts at location min(*µ*_*a*_, log 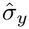) and scale 0.5, with the scale bounded above by 5.

### Supplementary Note 2: Initialization of the cohort model

Beyond the terms of Eqs. (16) and (17), the cohort model can include two more: a per-session deviation of the baseline rate, 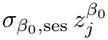, and a residual per-session deviation of the drug effect, 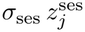. A term that the design cannot identify absorbs the variance of the term at the level below, so we include a term only when the assignment of sessions to animals and conditions identifies it. The animal-by-condition interaction *σ*_an*×d*_ needs at least one animal recorded under more than one condition, the residual *σ*_ses_ needs a repeated animal-by-condition cell, and the session term of the baseline 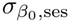 needs an animal recorded more than once. In the analyses reported here, the fitted terms were the animal terms and the animal-by-condition interaction in the crossed design, and the animal terms alone in the incomplete design. The simulated cohorts also include a per-session deviation of the baseline rate (standard deviation 0.20), which the fitted model does not represent.

The width of a drug contrast is set by the animal-by-condition interaction, which reaches *µ*_*d*_ through the non-centered effects **z**. A variational family that factorizes over **z** ignores this dependence, and with a Gaussian block over the population terms Θ alone the intervals were narrower than the spread of the estimate across simulated cohorts. The same held for the tonic contrast. On simulated cohorts without a tonic difference, a family that factorized the per-session tonic deviations reported intervals several times narrower than this spread, and including those deviations in the block brought the width of the interval in line with it without changing the rate contrast.

We initialize the cohort model recording by recording, in the same way as the single-trace model (Supplementary Note 1): each session gets its own tonic pre-fit and matched filter, from which its window scores start, and the per-condition means of the tonic response start at the least-squares pre-fits of the sessions of that condition. The joint Gaussian of Eq. (20) starts with 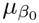 at the mean over sessions of the matched-filter estimate 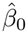, the *µ*_*d*_ at zero, the logarithm of each variance component at the location of its prior, the per-condition tonic means at the pre-fits, and the non-centered effects and tonic deviations at zero. Its initial scales are the prior scales for 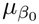 and the *µ*_*d*_, 0.1 for the variance components and 0.5 for the non-centered effects.

### Supplementary Note 3: Calibration of simulated recordings

We calibrated the simulated recordings against the experimental ones on the quantities that can be measured in both. For this calibration, the recordings were preprocessed with an asymmetric-least-squares smoothness of 10^4^ *×f*_*s*_, in place of the 10^6^*× f*_*s*_ used in the analyses (Methods). We measured the kernel and the noise on one session per animal, within the analysis window and at the sampling rate of the recordings. For the kernel, we fitted the bi-gamma form of the model by least squares to the peak-triggered average of 300 isolated transients. For the noise, we estimated the autoregressive coefficients of each session by the Yule–Walker method and took their mean, [1.432, *−*0.4464]. The leading coefficient ranged from 0.71 to 1.67 across the 10 sessions, so the simulated noise has the average color of the cohort and not that of a particular animal. We also tried adding an independent white measurement-noise term to the autoregressive process. Its estimated variance was below 10^*−*5^ of the total, and it worsened the Akaike information criterion in every animal, so we did not include it.

We set the noise scale by matching the ratio of transient height to local noise between simulated and experimental traces. We measured this ratio with the same procedure on both and obtained 4.4 *±*0.2 across the recordings and 4.2 in simulations at *σ*_*y*_ = 3. The same procedure is needed on both because the detector reports only peaks more than three local standard deviations above the noise. It misses the smaller transients, so the median of the detected peaks is higher than the median of all simulated amplitudes.

The event rate could not be calibrated in the same way, because on correlated noise a prominence detector returns peaks even in the absence of events. A trace simulated without events yields 12 detected peaks per minute, and a trace with 159 events per minute yields 9. At the baseline rate of the benchmark (*β*_0_ = *−*2.3, 6.0 events per minute), we detect 11 peaks per minute, compared with 13 *±*4 in the recordings. At these noise levels the detected peak rate is therefore mostly a property of the noise, so we set *β*_0_ instead of fitting it.

The benchmark recordings have the length, sampling rate and injection time of the experimental recordings (40 min at 48 Hz, injection at 10 min). Events occur at a baseline rate 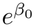 with *β*_0_ = *−*2.3 (6.0 events per minute), and the drug multiplies this rate by 2. Amplitudes follow LogNormal(1.77, 0.75^2^), with a median of 5.9 in arbitrary units. Their spread equals that of the fitted model, *σ*_*r*_, and their median sets the SNR at each noise level. The tonic response has height *P* = 35.1, so that its height relative to the noise is the same as in the cohort benchmark. Its rate constants are *k*_rel_ = 0.005 and *k*_cl_ = 0.0009 s^*−*1^, which place its peak about 7 min after the injection. We did not simulate drift. The kernel and the autoregressive noise are those calibrated above, and we compare the simulated and experimental spectra in Supplementary Fig. 1. The spectral distance shown in that figure does not distinguish first-from second-order noise, because each misses a different end of the spectrum. We therefore chose the order by the Akaike information criterion (Methods). With *p* = 2, the simulations contain 91% of the power of the recordings in the band of the tonic response (0.03–0.3 Hz) and 101% in the band of the transients (0.3–2 Hz). They contain 50% of the power below 0.03 Hz and 39% between 2 and 20 Hz.

### Supplementary Note 4: Alignment of detections to transient onsets

GuPPy and z-peaks report the peak of a transient, whereas the model reports its onset, which with this kernel comes 479 ms before the peak. Matching all methods to the true onsets without correcting for this would penalize GuPPy and z-peaks for a different definition of event time rather than for an error. We therefore estimated the systematic offset of each detector as the median signed distance from its detections to the nearest true onset within 979 ms (the time to peak of the kernel plus 0.5 s), and subtracted it from its detection times before matching. Across the 90 cells of the benchmark, the offset was 438 ms for GuPPy (range 417–458 ms) and 458 ms for z-peaks (range 417–458 ms), and 0 ms for the model, which estimates onsets. All GuPPy and z-peaks results in this paper include this correction, which on a recording at high SNR raised *F*_1_ from 0.128 to 0.697 for GuPPy and from 0.082 to 0.681 for z-peaks. The offsets are estimated on the same recordings to which they are applied, which favors GuPPy and z-peaks, since the model needs no correction.

## Supplementary Figures

**Supplementary Figure 1.**
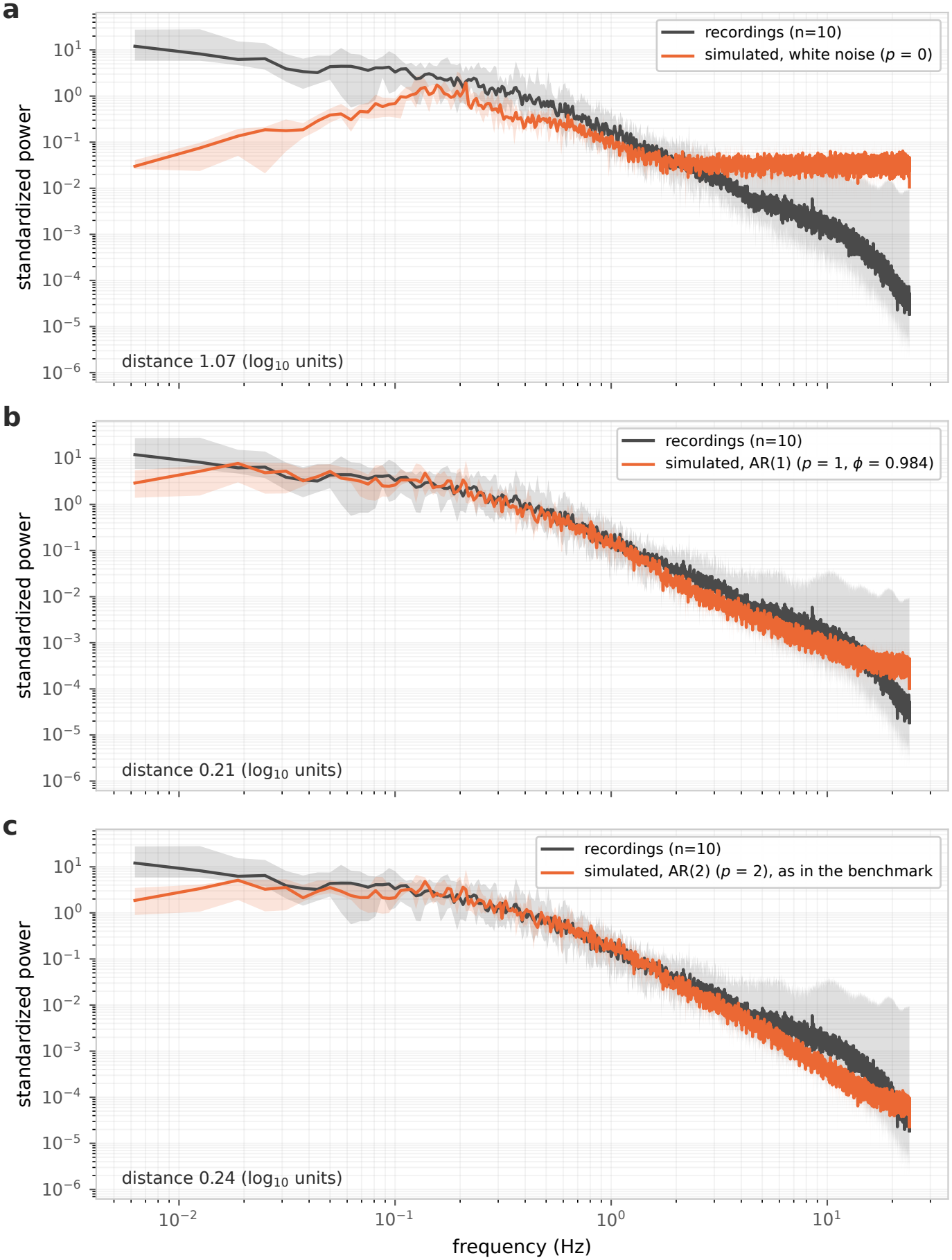
Power spectra of simulated and experimental recordings. Welch spectra of the 10 pre-injection windows of the recordings (grey) and of simulated recordings at *σ*_*y*_ = 3 (orange) with **a**, white noise (*p* = 0), **b**, first-order autoregressive noise (*p* = 1, *ϕ* = 0.984) and **c**, second-order autoregressive noise (*p* = 2; the benchmark setting). Each recording was divided by its robust scale before the transform. Lines, medians; bands, range across recordings. The number in each panel is the root-mean-square base-10 log ratio of the median spectra over 40 logarithmically spaced frequency bins from 0.01 Hz to the Nyquist frequency (1.07, 0.21 and 0.24 for *p* = 0, 1 and 2).

**Supplementary Figure 2.**
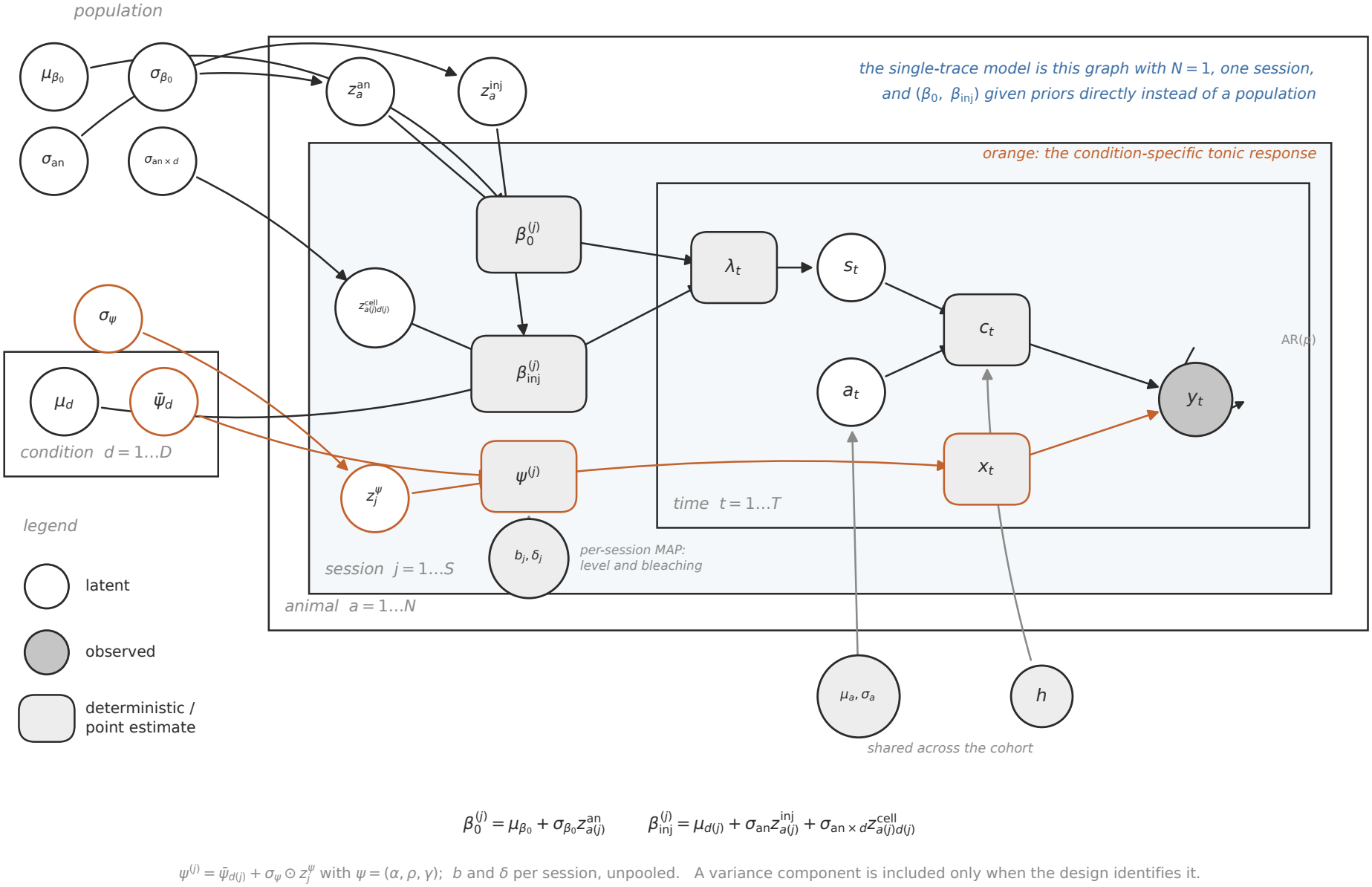
Graphical model of the cohort model. Conditional structure of Eqs. (16), (17) and (19) in plate notation. Nodes inside a plate are replicated over it. Animal effects lie outside the session plate, and the kernel and amplitude parameters, shared across the cohort, lie outside all plates. The single-trace model is the same graph with one animal and one session, and with priors placed directly on *β*_0_ and *β*_inj_. Orange, condition-specific tonic response; self-loop on *y*_*t*_, autoregressive emission, under which the observations within a recording are not conditionally independent.

**Supplementary Figure 3.**
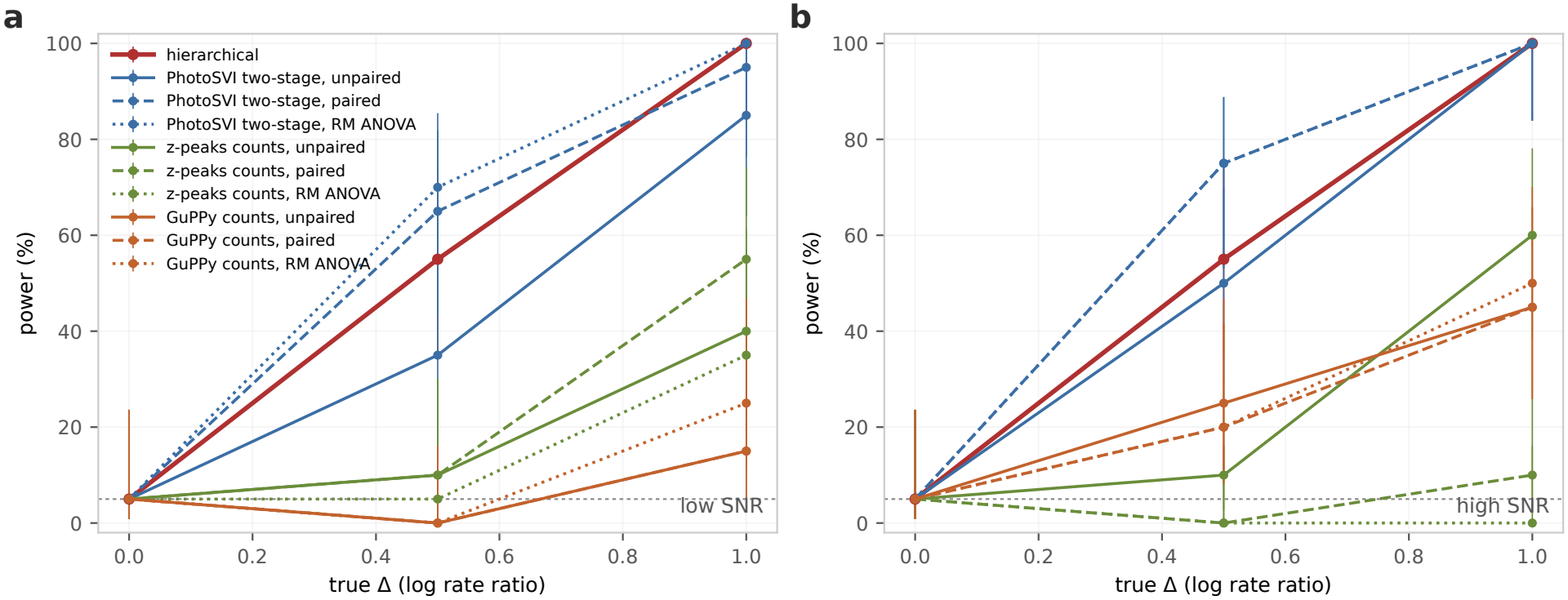
Power to detect a drug effect at low and high SNR. As Fig. 3a, for Δ*∈* {0, 0.5, 1} and 20 cohorts per effect size. **a**, SNR 1.5. **b**, SNR 2.9.

**Supplementary Figure 4.**
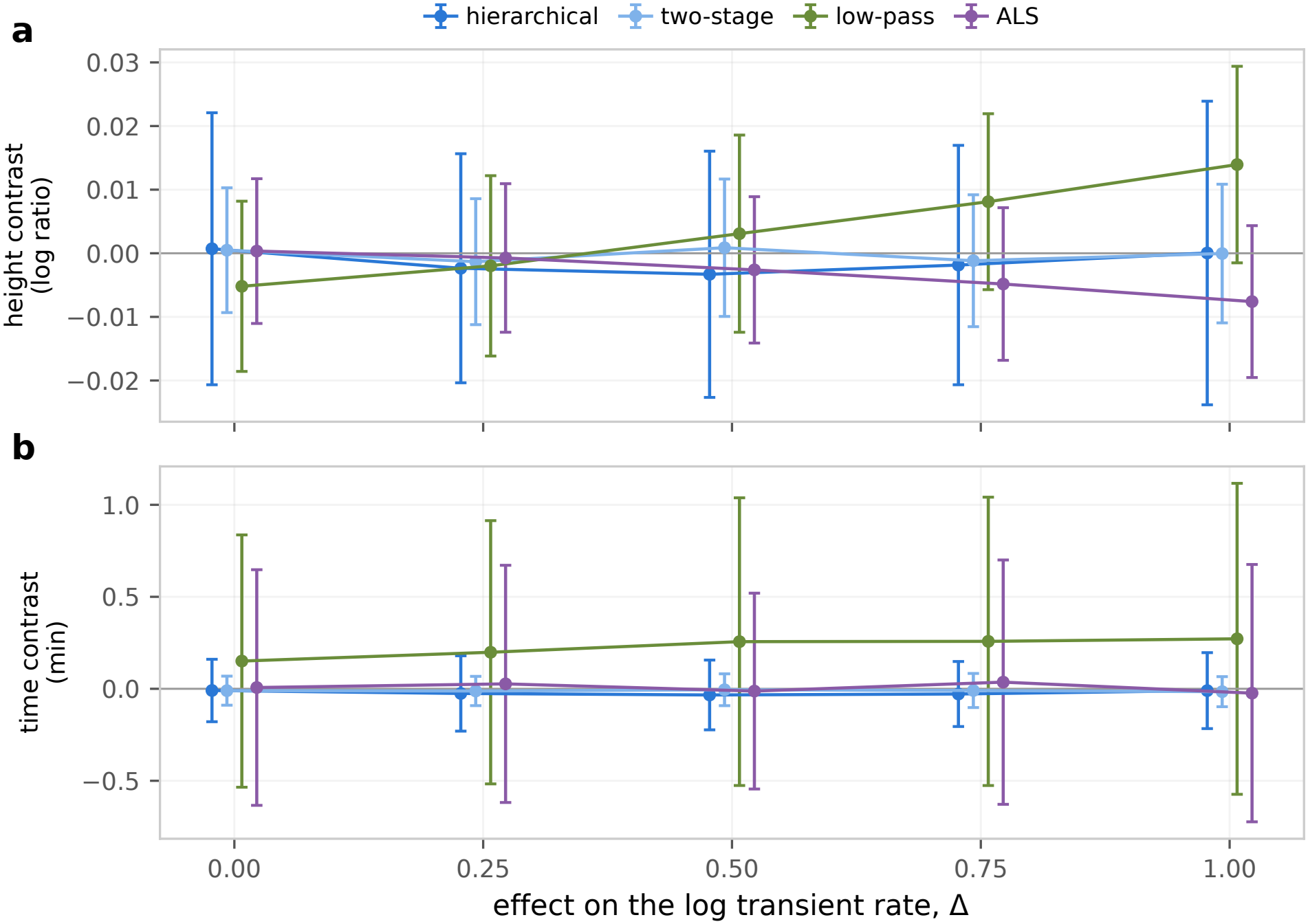
The tonic contrast in the cohort benchmark. Simulated cohorts at medium SNR (2.0; 50 per effect size), in which the treatment changes the log transient rate by Δ and the tonic response is the same in both conditions, so that the true tonic contrast is zero. **a**, Contrast between conditions in the height of the tonic peak (log ratio, treated minus vehicle). **b**, Contrast in the time of the tonic peak (minutes, treated minus vehicle). Hierarchical, posterior means of the cohort model; two-stage, single-trace PhotoSVI fits; low-pass and asymmetric least squares (ALS), the slow components of the same recordings (Methods). Two-stage, low-pass and ALS estimates are paired within animal. Mean *±* s.d. over cohorts.

**Supplementary Figure 5.**
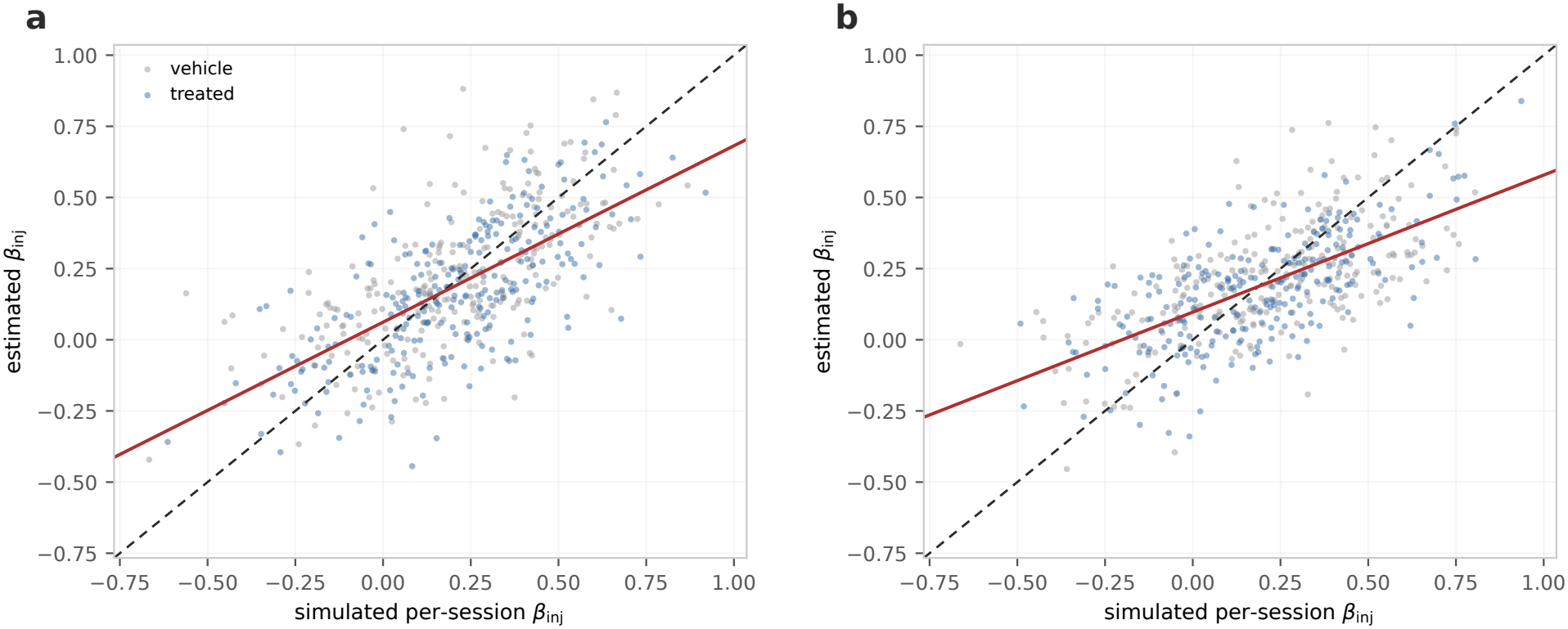
Shrinkage of the per-session estimates in the crossed and incomplete designs. Null cohorts (Δ = 0; 50 per design) at medium SNR (2.0), with the same number of recordings in both designs. Per-session estimates of *β*_inj_ against their simulated values; dashed line, identity; red line, least-squares fit. **a**, Crossed design, 5 animals recorded under both conditions (slope 0.62). **b**, Incomplete design, 10 animals recorded under one condition each (slope 0.48).

## Supplementary Tables

**Supplementary Table 1.**
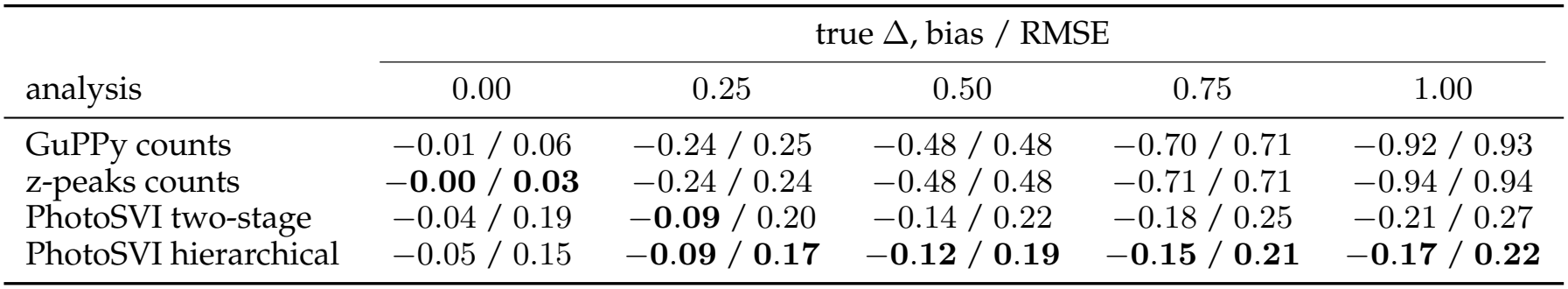
Accuracy of the estimated drug effect in simulated cohorts. Bias and root-mean-square error (RMSE) of the estimate of Δ given by each analysis, for the same cohorts and fits as in Fig. 3a. Paired and unpaired *t*-tests use the same estimate. At Δ = 0, estimates shrunk toward zero have small error by construction.

| analysis | true Δ, bias / RMSE |  |  |  |  |
|---|---|---|---|---|---|
|  | 0.00 | 0.25 | 0.50 | 0.75 | 1.00 |
| GuPPy counts | −0.01 / 0.06 | −0.24 / 0.25 | −0.48 / 0.48 | −0.70 / 0.71 | −0.92 / 0.93 |
| z-peaks counts | **−0.00** / **0.03** | −0.24 / 0.24 | −0.48 / 0.48 | −0.71 / 0.71 | −0.94 / 0.94 |
| PhotoSVI two-stage | −0.04 / 0.19 | **−0.09** / 0.20 | −0.14 / 0.22 | −0.18 / 0.25 | −0.21 / 0.27 |
| PhotoSVI hierarchical | −0.05 / 0.15 | **−0.09** / **0.17** | **−0.12** / **0.19** | **−0.15** / **0.21** | **−0.17** / **0.22** |

**Supplementary Table 2.** Accuracy of the estimated drug effect at low SNR. As Supplementary Table 1, at an SNR of 1.5.

|  | true Δ, bias / RMSE |  |  |
|---|---|---|---|
| analysis | 0.00 | 0.50 | 1.00 |
| GuPPy counts | −0.02 / 0.06 | −0.49 / 0.49 | −0.95 / 0.95 |
| z-peaks counts | **−0.00 / 0.03** | −0.49 / 0.49 | −0.96 / 0.96 |
| PhotoSVI two-stage | −0.04 / 0.20 | −0.19 / 0.27 | −0.29 / 0.35 |
| PhotoSVI hierarchical | −0.08 / 0.16 | **−0.18 / 0.23** | **−0.24 / 0.28** |

**Supplementary Table 3.**
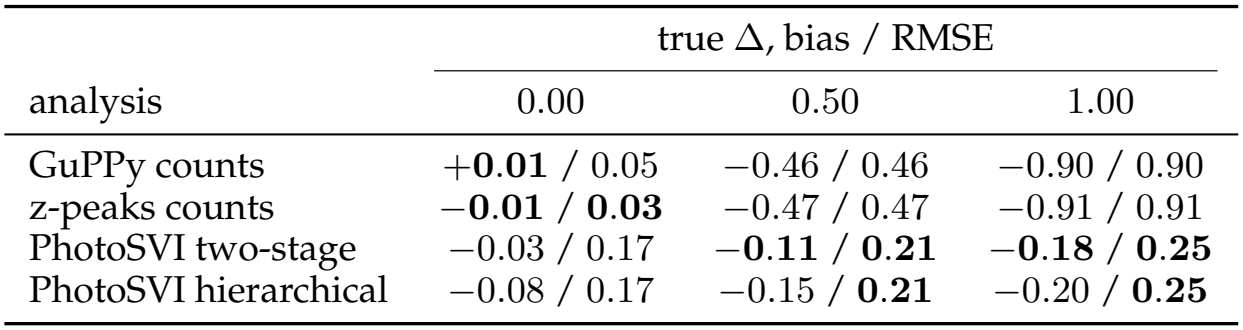
Accuracy of the estimated drug effect at high SNR. As Supplementary Table 1, at an SNR of 2.9.

|  | true Δ, bias / RMSE |  |  |
|---|---|---|---|
| analysis | 0.00 | 0.50 | 1.00 |
| GuPPy counts | +**0.01** / 0.05 | −0.46 / 0.46 | −0.90 / 0.90 |
| z-peaks counts | −**0.01** / **0.03** | −0.47 / 0.47 | −0.91 / 0.91 |
| PhotoSVI two-stage | −0.03 / 0.17 | −**0.11** / **0.21** | −**0.18** / **0.25** |
| PhotoSVI hierarchical | −0.08 / 0.17 | −0.15 / **0.21** | −0.20 / **0.25** |

**Supplementary Table 4.** The effect of the pretreatment in the opioid cohort, by analysis. Ratio of the morphine-induced increase in transient rate after RO64-6198 to that after vehicle. Hierarchical, the cohort-model estimate reported in the main text. Two-stage, each session’s ratio of post-to pre-injection rate, from the expected number of events under PhotoSVI or from the detections of GuPPy and z-peaks at their default settings, compared between pretreatments by a paired *t*-test over the animals recorded under both or an unpaired *t*-test over all sessions (Methods).

|  | paired within animal |  | unpaired |  |
|---|---|---|---|---|
| analysis | ratio [90% interval] | *p* | ratio | *p* |
| PhotoSVI hierarchical | 1.57 [1.13, 2.17] | – | – | – |
| PhotoSVI two-stage | 1.51 [1.28, 1.78] | 0.004 | 1.39 | 0.23 |
| z-peaks counts | 1.26 [1.05, 1.51] | 0.055 | 1.23 | 0.21 |
| GuPPy counts | 1.11 [0.95, 1.30] | 0.220 | 1.14 | 0.29 |

**Supplementary Table 5.**
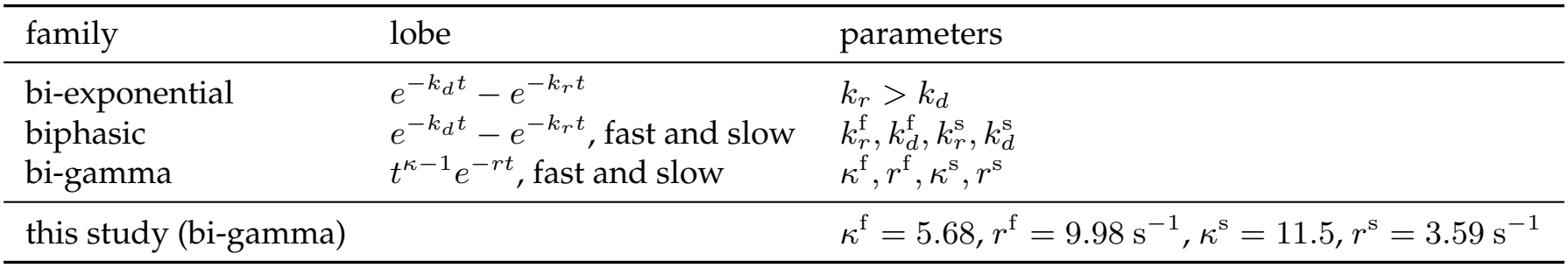
Sensor kernel families implemented in PhotoSVI. Biphasic and bi-gamma kernels are the difference of a fast and a slow lobe, weighted so that the kernel has zero area, ∑_*k*_ *h*_*k*_ = 0. The bi-exponential kernel is a single lobe made zero-area by subtracting its mean over the support. All kernels are normalized to unit peak; *t*, time since the event onset; *κ*, shape; *r*, rate. The kernel used in this study is the bi-gamma kernel fitted to the peak-triggered average of 300 isolated transients of the experimental recordings (dLight1.3b), truncated at 400 bins (8.3 s).

| family | lobe | parameters |
|---|---|---|
| bi-exponential | e<sup>−k<sub>d</sub>t</sup> − e<sup>−k<sub>r</sub>t</sup> | k<sub>r</sub> > k<sub>d</sub> |
| biphasic | e<sup>−k<sub>d</sub>t</sup> − e<sup>−k<sub>r</sub>t</sup>, fast and slow | k<sub>r</sub><sup>f</sup>, k<sub>d</sub><sup>f</sup>, k<sub>r</sub><sup>s</sup>, k<sub>d</sub><sup>s</sup> |
| bi-gamma | t<sup>κ−1</sup>e<sup>−rt</sup>, fast and slow | κ<sup>f</sup>, r<sup>f</sup>, κ<sup>s</sup>, r<sup>s</sup> |
| this study (bi-gamma) |  | κ<sup>f</sup> = 5.68, r<sup>f</sup> = 9.98 s<sup>−1</sup>, κ<sup>s</sup> = 11.5, r<sup>s</sup> = 3.59 s<sup>−1</sup> |

**Supplementary Table 6.**
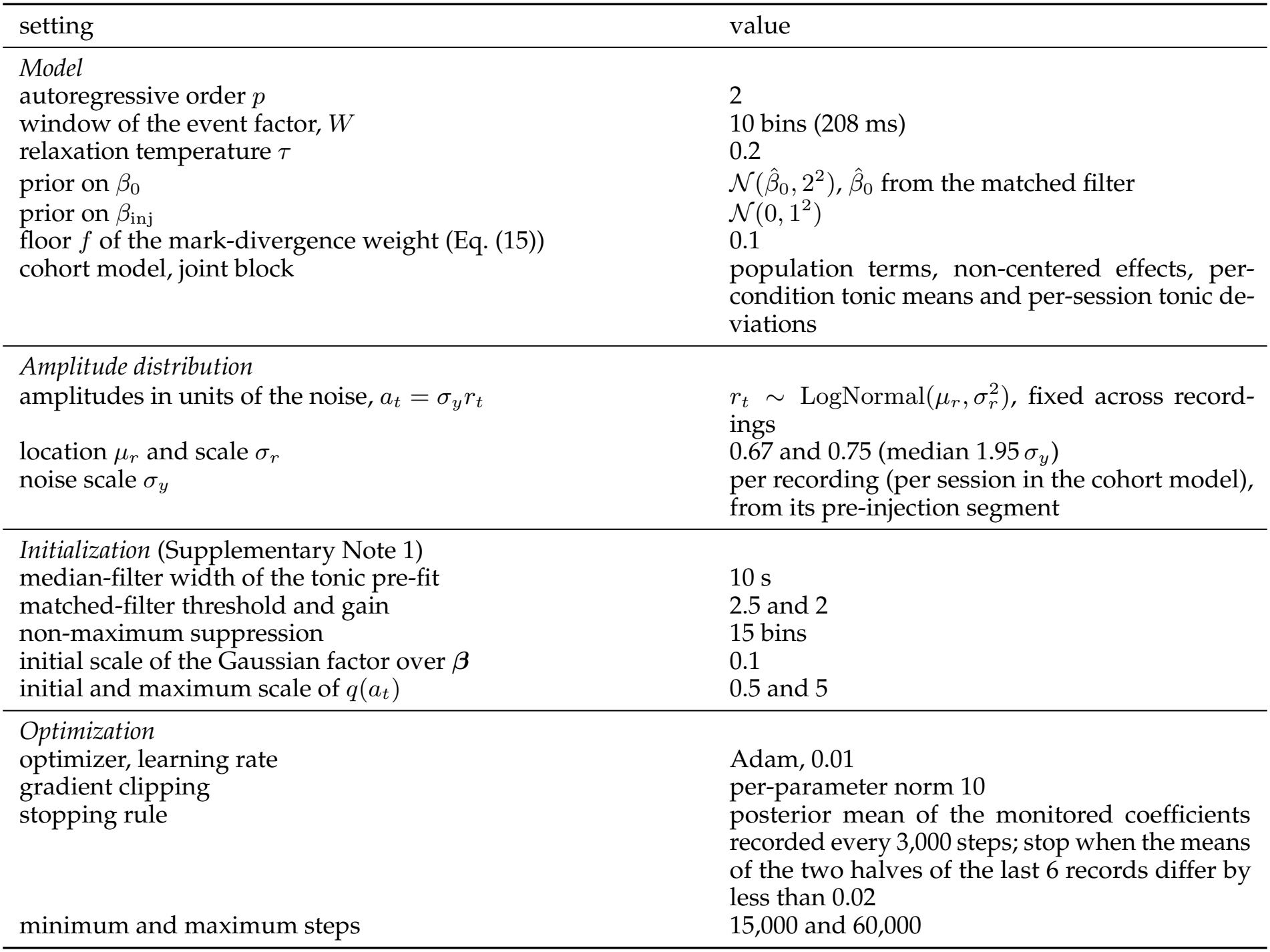
Fixed settings of the model and its fit. The same settings are used in every fit, single-trace and cohort, except where the cohort model is indicated.

| setting | value |
|---|---|
| *Model* |  |
| autoregressive order *p* | 2 |
| window of the event factor, *W* | 10 bins (208 ms) |
| relaxation temperature τ | 0.2 |
| prior on β<sub>0</sub> | $\mathcal{N}(\hat{\beta}_0, 2^2)$, $\hat{\beta}_0$ from the matched filter |
| prior on β<sub>inj</sub> | $\mathcal{N}(0, 1^2)$ |
| floor *f* of the mark-divergence weight (Eq. (15)) | 0.1 |
| cohort model, joint block | population terms, non-centered effects, per-condition tonic means and per-session tonic de-viations |
| *Amplitude distribution* |  |
| amplitudes in units of the noise, *a*<sub>*t*</sub> = σ<sub>*y*</sub>*r*<sub>*t*</sub> | *r*<sub>*t*</sub> ∼ LogNormal(μ<sub>*r*</sub>, σ<sub>*r*</sub><sup>2</sup>), fixed across record-ings |
| location μ<sub>*r*</sub> and scale σ<sub>*r*</sub> | 0.67 and 0.75 (median 1.95 σ<sub>*y*</sub>) |
| noise scale σ<sub>*y*</sub> | per recording (per session in the cohort model), from its pre-injection segment |
| *Initialization* (Supplementary Note 1) |  |
| median-filter width of the tonic pre-fit | 10 s |
| matched-filter threshold and gain | 2.5 and 2 |
| non-maximum suppression | 15 bins |
| initial scale of the Gaussian factor over **β** | 0.1 |
| initial and maximum scale of *q*(*a*<sub>*t*</sub>) | 0.5 and 5 |
| *Optimization* |  |
| optimizer, learning rate | Adam, 0.01 |
| gradient clipping | per-parameter norm 10 |
| stopping rule | posterior mean of the monitored coefficients recorded every 3,000 steps; stop when the means of the two halves of the last 6 records differ by less than 0.02 |
| minimum and maximum steps | 15,000 and 60,000 |

